# An interdependent Metabolic and Genetic Network shows emergent properties *in vitro*

**DOI:** 10.1101/2023.11.26.568713

**Authors:** Simone Giaveri, Nitin Bohra, Christoph Diehl, Martine Ballinger, Nicole Paczia, Timo Glatter, Tobias J. Erb

## Abstract

A hallmark of all living organisms is their ability for self-regeneration which requires a tight integration of metabolic and genetic networks. Here we constructed a metabolic and genetic linked in vitro network (MGLN) that shows life-like behavior outside of a cellular context and generates its own building blocks from non-living matter. To this end, we integrated the metabolism of the crotonyl-CoA/ethyl-malonyl-CoA/hydroxybutyryl-CoA (CETCH) cycle with cell-free protein synthesis using recombinant elements (PURE). We demonstrate that the MGLN produces the essential amino acid glycine from inorganic carbon (CO_2_), and incorporates it into target proteins following DNA-encoded instructions. By programming genetically encoded response into metabolic networks our work opens new avenues for the development of advanced biomimetic systems with emergent properties, including decision-making, self-regeneration and evolution.

## Main Text

All living systems show emergent properties such as self-organization, regeneration and repair. Cells solved this challenge by a tight integration of metabolic and genetic networks (*1*, *2*). Genetic networks encode the components of metabolic networks that in turn provide the energy and building blocks to execute the genetic programs for self-regulation and -regeneration (*3*). Constructing systems that show similar functionalities is one of the milestones on the way to construct synthetic cells, but also key to develop biotechnological systems that are capable of decision-making, self-optimization and Darwinian evolution (*4*).

Recent efforts in synthetic biology pioneered cell-free transcription-translation (TX-TL) systems that are able to process genetic programs (*5*, *6*) and (partially) self-regenerate when supplemented with instructions, energy, and building blocks (*7*, *8*). At the same time, several complex metabolic networks have been successfully reconstructed outside of a cellular context (*9–16*). These *in vitro* networks were combined with energy modules (*17*, *18*) to produce different metabolites, including organic acids and more complex molecules from simple precursors, such as glucose or CO_2_ (*19*).

Yet, so far, such genetic and metabolic *in vitro* systems have not been integrated with each other to achieve higher-order functionality. Here, we aimed at developing a metabolically and genetically linked network (MGLN), in which both layers are recursively communicating with each other. This link is achieved through an essential amino acid (glycine), which is autonomously synthesized from inorganic carbon (CO_2_) through the metabolic network, controlling the output of the genetic network, which itself feeds back to the metabolic layer through DNA-encoded instructions.

To realize such a MGLN network, we sought to combine a 16-enzyme *in vitro* CO_2_-fixation system, the CETCH cycle (*9*), with a cell-free TX-TL system (PURE) for protein production (*20*). The CETCH cycle converts CO_2_ into glyoxylate, which we aimed to turn into glycine and incorporate into target proteins through the TX-TL machinery of the PURE system. With these proteins, we further envisioned to establish recursive feed-forward loops to the metabolic network that would allow the production of key metabolic enzymes of the CETCH cycle (Fig. 1).

**Fig. 1.**
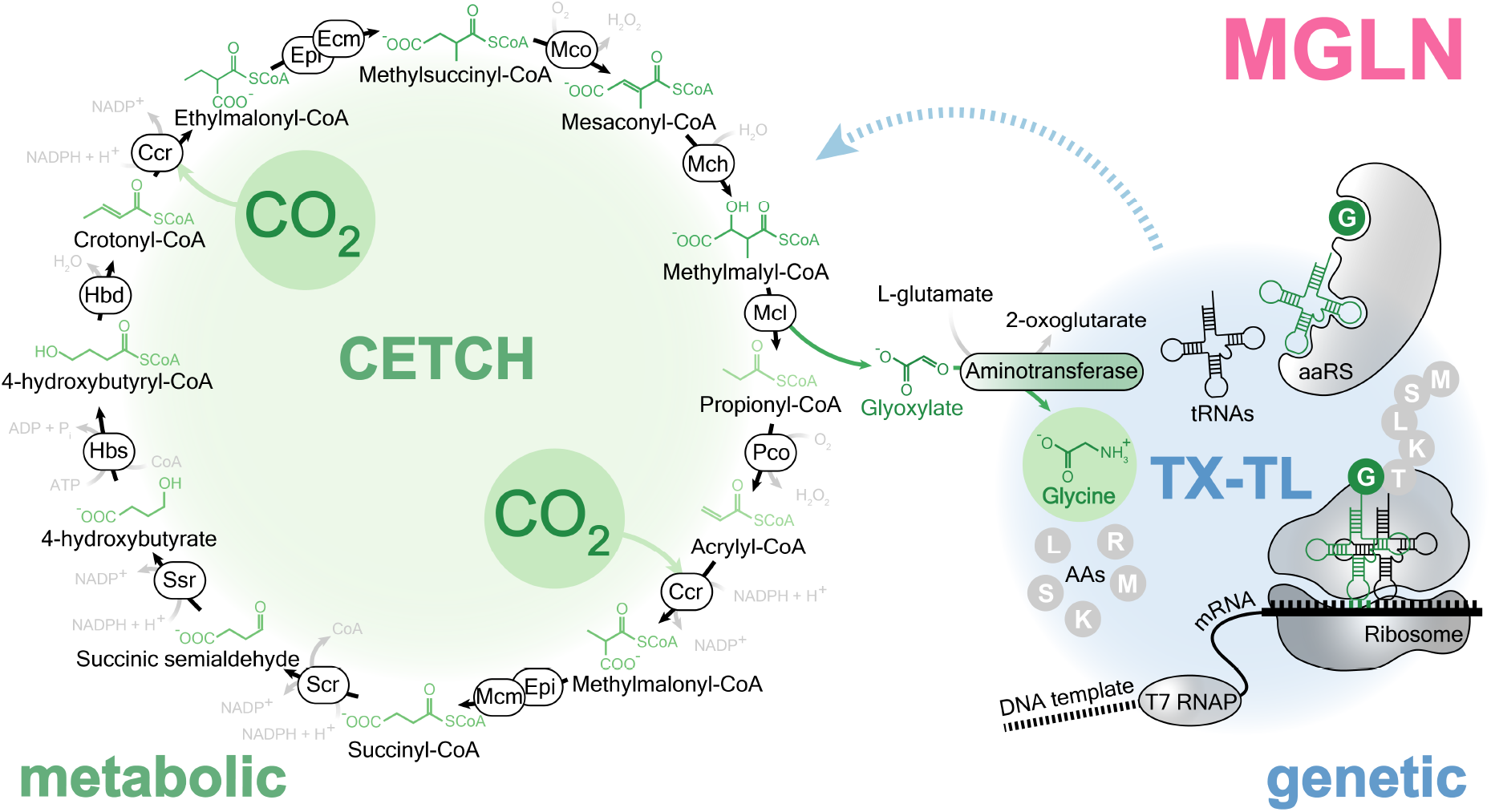
Experimental design of a MGLN. Schematic of the topology of the MGLN described in this study, where the metabolic layer of the CETCH cycle (green) is integrated with the genetic layer of TX-TL (blue) by the transamination of the product of CO_2_ fixation (glyoxylate) into an amino acid (glycine); glycine is used in turn by TX-TL to produce enzymes key for CO_2_ fixation. See Table S1 for enzyme abbreviations.

To construct such a system, we first established a PURE TX-TL mix lacking glycine (Δgly-PURE TX-TL) in which externally added glycine is required for protein production. We provided this Δgly-PURE TX-TL system with a DNA template encoding for the green fluorescent protein EGFP (*21*) and supplemented it with different concentrations of glycine. Fluorescence read-out showed a linear dependency between 10 μM and 150 μM glycine (Fig. 2a), confirming that the concentration of glycine directly controls EGFP production in the Δgly-PURE system.

**Fig. 2.**
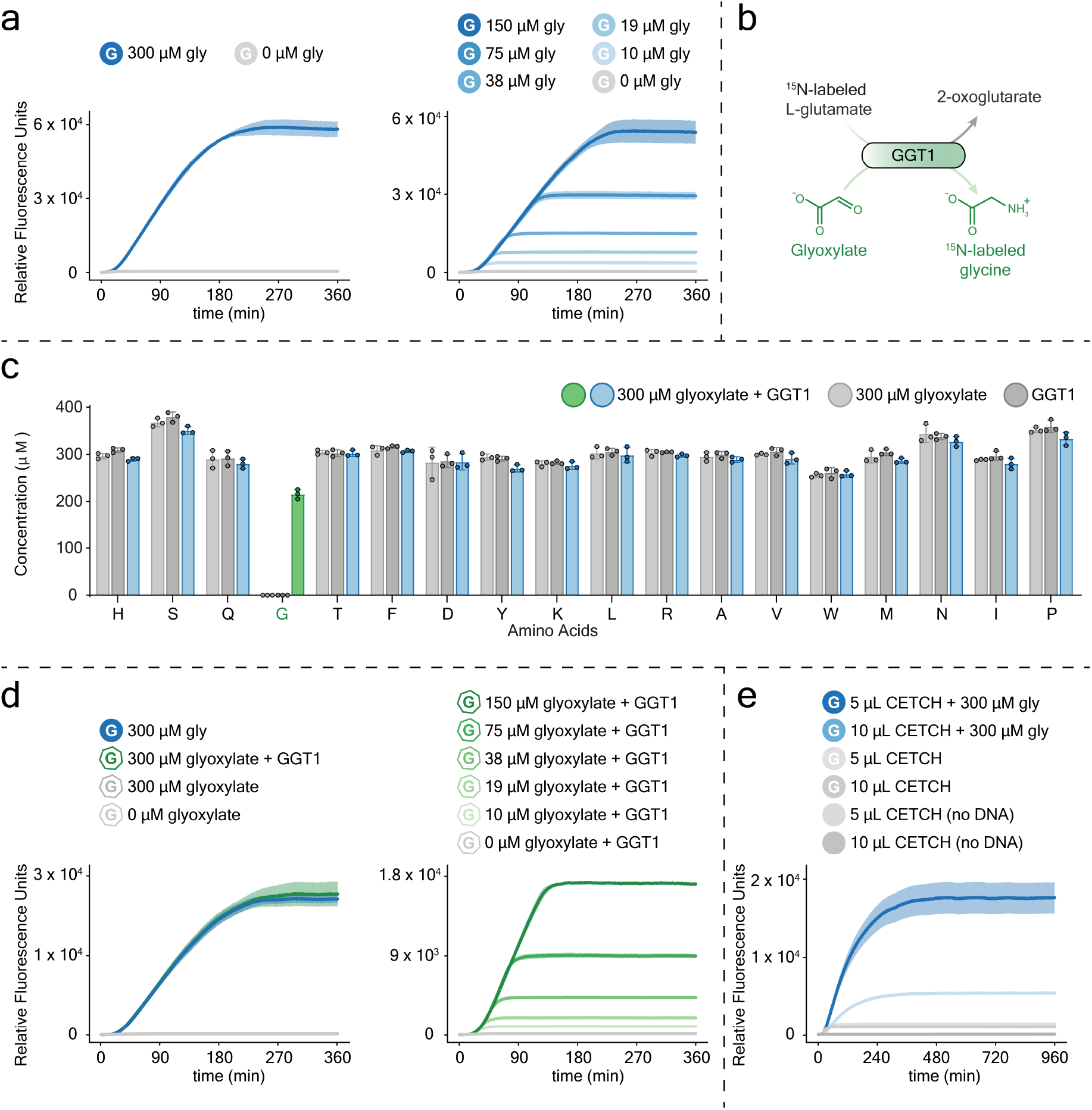
Control of cell-free TX-TL with glycine, or glyoxylate. (**a**) Fluorescent signal resulting from the cell-free expression of *EGFP* template in Δgly-PURE TX-TL system with different concentrations of externally added glycine (blue curves). Negative controls without glycine were included (grey curves). TX–TL reactions were incubated at 37 °C into the microplate reader, with gain = 100. (**b**) Scheme of the transamination reaction of glyoxylate into ^15^N-glycine by ^15^N-glutamate, and GGT1 aminotransferase from *Arabidopsis thaliana*. (**c**) Bar graph representing the amino acid analysis by HPLC-MS/MS of the result of the transamination reaction sketched in (b), performed in presence of GGT1, glyoxylate, ^15^N-glutamate, and 19 proteinogenic amino acids (glycine omitted), for 15 min (colored bars, ^15^N-glycine is highlighted in green), at 30 °C. Negative control assays without GGT1 (light grey bars), or glyoxylate (dark grey bars) were included. Comprehensive data of the amino acid analysis are reported in Figs. S2-6. (**d**) Fluorescent signal resulting from the cell-free expression of *EGFP* template in Δgly-PURE TX-TL system with GGT1 and different concentrations of externally added glyoxylate (green curves). Negative controls without glyoxylate, or without GGT1 were performed (grey curves). A reference control with glycine (blue curve) was included. TX–TL reactions were incubated at 37 °C into the microplate reader, with gain = 90. (**e**) Fluorescent signal resulting from the cell-free expression of *EGFP* template in Δgly-PURE TX-TL system supplemented with glycine, in presence of different concentrations of CETCH components added into 25 μl TX-TL reactions, without propionyl-CoA (blue curves). Negative controls without propionyl-CoA and glycine, or without propionyl-CoA, and DNA templates (grey curves) were performed. TX–TL reactions were incubated at 37 °C into the microplate reader, with gain = 90. All TX–TL reactions were run in triplicates; expression curves represent the statistical mean of the results at any acquisition time; shadows represent the standard deviation of the same data. Bar-plots of the statistical mean of the results (triplicates) are shown; error bars represent the standard deviation of the same data.

Next, we tested whether glycine could be produced *in situ* in the Δgly-PURE system from externally added glyoxylate (the product of the CETCH cycle). We sought to employ a transamination of glyoxylate by accessing the large pool of potassium glutamate (100 mM) of the PURE system and using glutamate-glyoxylate amino-transferase (GGT1) from *Arabidopsis thaliana* (*22*) or the promiscuous aspartate-glyoxylate aminotransferase (BhcA) from *Paracoccus denitrificans* (*23*) as transaminating enzyme (Fig. 2b). We tested either aminotransferase in a reaction mixture with 100 μM pyridoxal phosphate (PLP), 300 μM glyoxylate, 100 mM Potassium ^15^N-glutamate and all 19 proteinogenic amino acids (at 300 μM each), and followed the formation of ^15^N-labelled and unlabeled glycine with high-performance liquid chromatography-mass spectrometry (HPLC-MS/MS). GGT1 produced 217 ± 15 μM ^15^N-glycine after 15 min, and only minor amounts of unlabeled glycine (Fig. 2c), demonstrating that glutamate served as exclusive amino donor for GGT1. In contrast, BhcA produced 201 ± 6 μM ^15^N-glycine and 40 ± 3 μM unlabeled glycine (Fig. S2), indicating that other amino acids (*i.e.*, aspartic acid, see Fig. S2) were used by BhcA. We chose to continue with GGT1 and integrated the transaminase into a Δgly-PURE TX-TL system with EGFP as fluorescent readout. Similar to the experiments with externally added glycine, we observed a linear dependence of EGFP production from glyoxylate in the range of 10-150 μM glyoxylate (Fig. 2d). These results showed that glyoxylate could indeed serve as link between the CETCH cycle and Δgly-PURE TX-TL system.

In the following, we tested the effect of coenzymes, cofactors, salts, and buffers of the CETCH cycle onto productivity of the Δgly-PURE TX-TL system. We decided for a version of the CETCH cycle (CETCH vA7) that was recently optimized by active learning (*24*), and added its components – except the starting substrate propionyl-CoA – to a Δgly-PURE TX-TL system with 300 µM glycine and EGFP as readout. When testing different ratios of the CETCH cycle and Δgly-PURE TX-TL, a 25 µl of PURE reaction mixture containing 5 μl of CETCH worked best (Fig. 2e), demonstrating that the Δgly-PURE system can be coupled with a 1:5 diluted CETCH cycle.

We next tested whether a 1:5 diluted CETCH cycle was able to produce sufficient glycine in presence of 100 mM potassium glutamate and GGT1. HPLC-MS/MS analysis showed that a diluted CETCH started with 200 μM propionyl-CoA produced 552 ± 27 μM glycine at 30 °C within 4 hours (Fig. 3a). This activity corresponded to a CO_2_-fixation efficiency 17_CO2_ of 5.5 ± 0.3 (17_CO2_ defined as CO_2_ equivalents fixed per acceptor molecule), which is comparable to earlier published versions of the CETCH cycle (*e.g.*, CETCH v5.4 (*9*)).

**Fig. 3.**
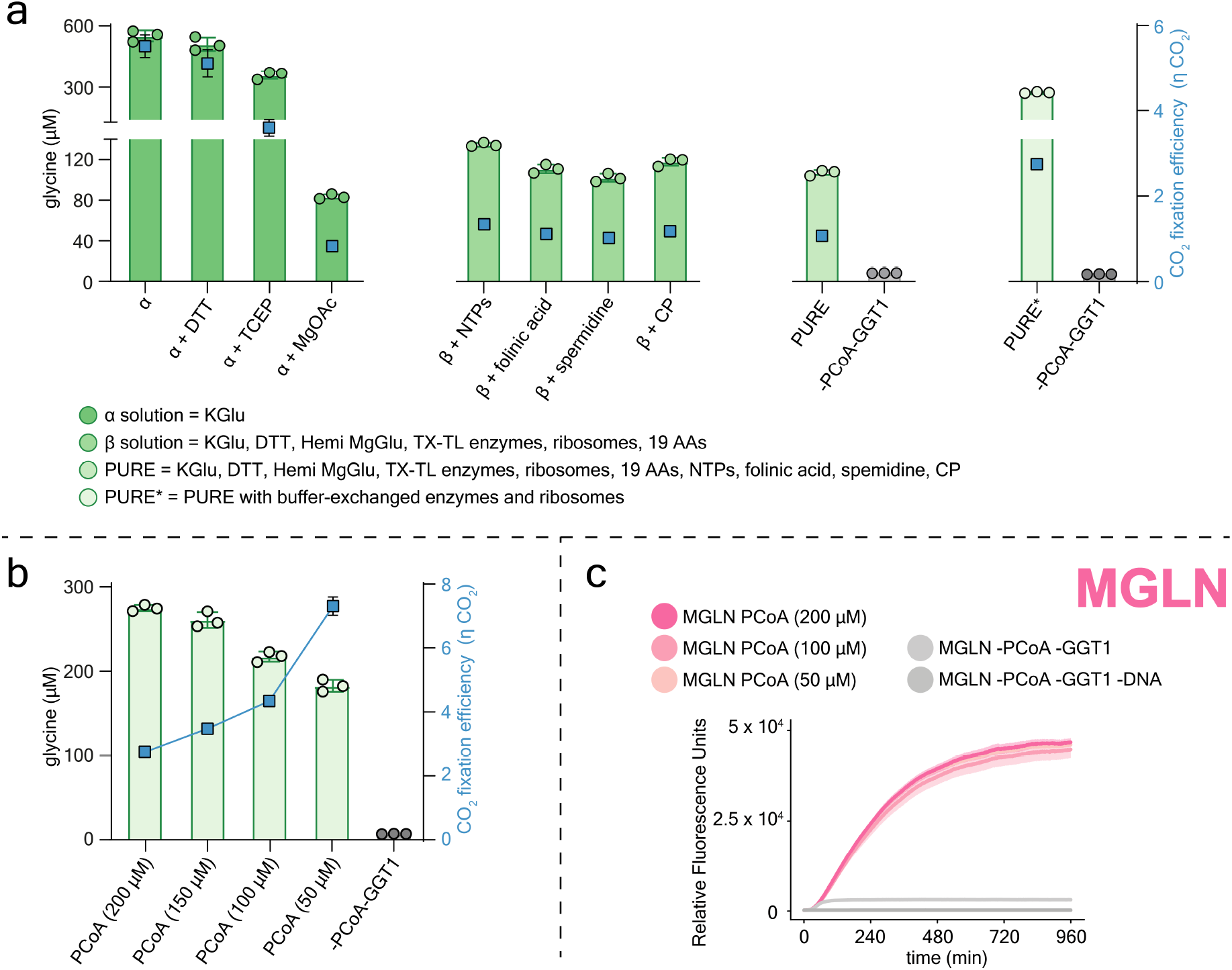
Continuous fixation of CO_2_ into glycine by the CETCH cycle supplemented with Δgly-PURE TX-TL system. (**a**) Left *y* axis: bar graph representing the absolute concentration of glycine, measured by HPLC-MS/MS, produced over the course of 4 h at 30 °C by the CETCH cycle started with 200 μM propionyl-CoA, and stepwise supplemented with the components of the Δgly-PURE TX-TL system (green bars). Negative controls without propionyl-CoA and GGT1 were included (grey bars). Right *y* axis: CO_2_-fixation efficiency 17_CO2_ of the CETCH cycle (blue squares). A detailed description of the reaction assembly is reported in Table S5. (**b**) Left *y* axis: bar graph representing the absolute concentration of glycine, measured by HPLC-MS/MS, produced over the course of 4 h at 30 °C by the CETCH cycle in presence of Δgly-PURE TX-TL system, and started with different concentrations of propionyl-CoA (green bars). Right *y* axis: CO_2_-fixation efficiency 17_CO2_ of the CETCH cycle (blue squares). (**c**) Fluorescent signal resulting from the cell-free expression of *EGFP* template in Δgly-PURE TX-TL system added with the CETCH cycle and different concentrations of propionyl-CoA (pink curves). Negative controls without propionyl-CoA and GGT1 (light grey curve), or without DNA templates (dark grey curve) were performed. TX–TL reactions were incubated at 30 °C into the microplate reader, with gain = 100. TX–TL reactions were run in triplicates; expression curves represent the statistical mean of the results at any acquisition time; shadows represent the standard deviation of the same data. Bar-plots of the statistical mean of the results (triplicates) are shown; error bars represent the standard deviation of the same data.

We continued by stepwise supplementing the 1:5 diluted CETCH cycle with additional components of the Δgly-PURE TX-TL system, to identify any factors that would affect 17_CO2_ (Fig. 3a). Upon addition of the reducing agents dithiothreitol (DTT), or Tris(2-carboxyethyl)phosphine (TCEP), 17_CO2_ decreased slightly to moderately (5.1 ± 0.3 for DTT; 3.6 ± 0.2 for TCEP). In contrast, supplementation with magnesium acetate (11.8 mM) severely decreased 17_CO2_ to 0.8 ± 0.02, indicating a strong negative interaction of system components. We hypothesized that acetate could serve as alternative substrate for 4-hydroxybutyryl-CoA synthetase (Hbs) in the CETCH, thus reducing flux through the cycle (*25*). To overcome this limitation, we replaced magnesium acetate with hemi-magnesium glutamate (*26*), and also removed any residual acetate in the ribosome and TX-TL enzyme fractions of the Δgly-PURE system via buffer exchange, which increased 17_CO2_ significantly to 2.8 ± 0.04, corresponding to the production of 275 ± 4 μM glycine within 4 hours (Fig. 3a).

Finally, we also investigated, how varying concentrations of the starting substrate propionyl-CoA would affect 17_CO2_ of the 1:5 diluted CETCH cycle. Decreasing propionyl-CoA concentrations from 200 to 50 μM, decreased total glycine production from 275 ± 4 μM to 183 ± 7 μM, but significantly increased the 17_CO2_ from 2.8 ± 0.04 to 7.3 ± 0.3 (Fig. 3b), indicating that the 1:5 diluted CETCH cycle turned more efficiently at lower propionyl-CoA concentrations.

Having optimized the working conditions of the metabolic (CETCH cycle) and genetic (Δgly-PURE TX-TL) layers independently and reciprocally, we aimed at operating both systems together at 30 °C *i.e.*, the optimized temperature for the CETCH cycle (Fig. S9). We combined the 16 enzymes of the CETCH cycle with the 36 enzymes of the Δgly-PURE TX-TL system, added GGT1, and supplemented the reaction mix with ribosomes, and all other coenzymes, cofactors, and salts required. We activated the metabolic layer with different concentrations of propionyl-CoA (50, 100, 200 µM, respectively) and the genetic layer with a DNA template encoding for EGFP. Under these conditions, EGFP production above background was only observed, when GGT1 was present (converting glyoxylate produced by the CETCH cycle into glycine used by Δgly-PURE TX-TL). Moreover, these results confirmed that 50 µM propionyl-CoA was sufficient and optimal for EGFP production (Fig. 3c), as higher concentrations of the starting substrate did not further improve the EGFP signal, which is in line with earlier experiments that showed EGFP signal saturation above 150 µM glycine (Fig. 2a, 2d). Altogether, these experiments showed the successful (self-)integration of the metabolic and genetic networks into one *in vitro* system.

Having successfully linked the metabolic and genetic layers (*i.e.*, the CETCH cycle and Δgly-PURE TX-TL) with each other, we next thought of establishing positive feedback between both systems. We envisioned an “autocatalytic” MGLN in which the metabolic and genetic activities are linked through a feed-forward loop. We designed a feed-forward loop based on GGT1, which we provided as DNA-based template to a PURE TX-TL system with an EGFP readout (Fig. S13). In the presence of 10 µM glycine, EGFP was produced at relevant concentrations, when external glyoxylate (300 µM) was provided (Fig. S10), or when the CETCH cycle was started with propionyl-CoA (Fig. S13).

Inspired by nature, we sought to further integrate our metabolic and genetic networks in order to achieve “self-regeneration” *in vitro*. To this end, we selected ethylmalonyl-CoA mutase (Ecm) from *Rhodobacter sphaeroides* that converts ethylmalonyl-CoA into methylsuccinyl-CoA (Fig. 1). We produced Ecm by TX-TL in the PURE system (Fig. S8), and verified the enzyme activity by co-expressing *Ecm* and *EGFP* templates in a Δgly-PURE TX-TL system supplemented with 100 μM ethylmalonyl-CoA, CETCH components (see Methods), and 10 μM glycine (Fig. S11). Next, we assembled a ΔEcm-MGLN, that is a cell-free system able to partially self-regenerate by producing the missing Ecm enzyme with the amino acid glycine, autonomously synthesized from CO_2_. Upon feeding ΔEcm-MGLN with the DNA sequences encoding for Ecm, (and EGFP), 50 μM propionyl CoA, as well as with minimal glycine concentrations (10 μM), the MGLN increased its CO_2_ fixation, in a feed-forward manner through its own metabolic activity (Figs. 4a, S14). Proteomic analysis using NaH^13^CO_3_ verified that the majority of glycine residues in Ecm, (and EGFP) were synthesized from CO_2_ (Fig. S15).

**Fig. 4.**
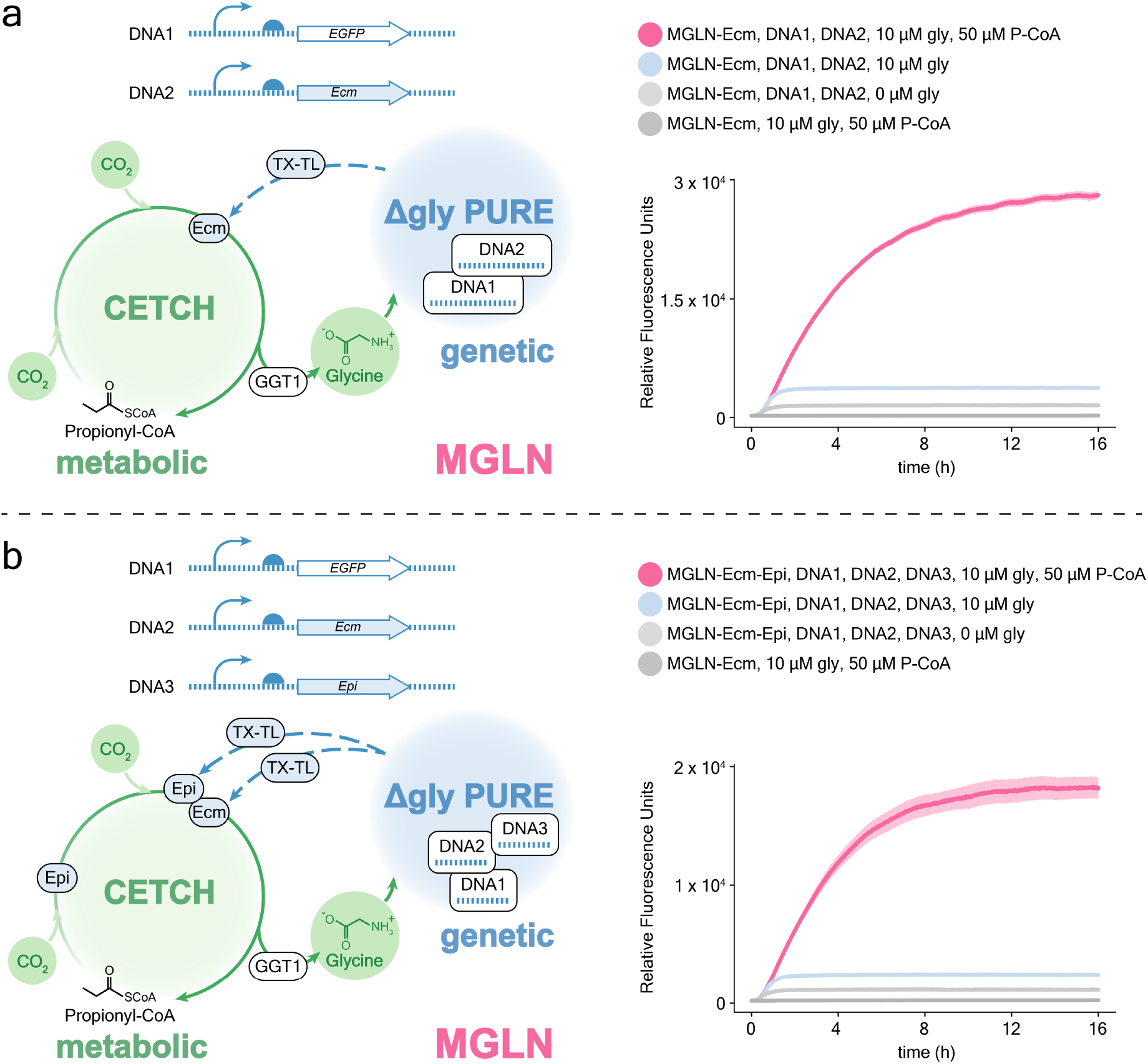
Self-regenerating MGLNs. (**a**) (Left) schematics of a ΔEcm-MGLN able to partially self-regenerate by producing the missing Ecm enzyme with the amino acid glycine, autonomously synthesized from CO_2_. (Right) fluorescent signal resulting from the cell-free expression of *EGFP* template in ΔEcm-MGLN added with *Ecm* template, propionyl-CoA, and 10 μM glycine concentration (pink curve). Negative controls without propionyl-CoA (blue curve), or without propionyl-CoA and glycine (light grey curve), or without DNA templates were included (dark grey curve). (**b**) (Left) schematics of a Δ(Ecm,Epi)-MGLN able to partially self-regenerate by producing the missing Ecm and Epi enzymes with the amino acid glycine, autonomously synthesized from CO_2_. (Right) fluorescent signal resulting from the cell-free expression of *EGFP* template in Δ(Ecm,Epi)-MGLN added with *Ecm* and *Epi* templates, propionyl-CoA, and 10 μM glycine concentration (pink curve). Negative controls without propionyl-CoA (blue curve), or without propionyl-CoA and glycine (light grey curve), or without DNA templates (dark grey curve) were performed. TX–TL reactions were incubated at 30 °C into the microplate reader, with gain = 100. TX–TL reactions were run in triplicates; expression curves represent the statistical mean of the results at any acquisition time; shadows represent the standard deviation of the same data. Bar-plots of the statistical mean of the results (triplicates) are shown; error bars represent the standard deviation of the same data.

We ultimately aimed at assembling a MGLN able to regenerate multiple enzymes of its metabolic layer. To this end, we selected the methylmalonyl-/ethylmalonyl-CoA epimerase (Epi) from *Rhodobacter sphaeroides* that converts ethylmalonyl-CoA into methylsuccinyl-CoA coupled with Ecm (Fig. 1). We produced Epi by cell-free TX-TL (Fig. S8), and verified the enzyme activity as described above (see Methods, and Fig. S12). Next, we assembled a Δ(Ecm,Epi)-MGLN that, upon providing the system with DNA sequences, propionyl CoA, and minimal concentrations of glycine, is able to self-regenerate by partially producing from CO_2_ the enzymes necessary for a core step of the CETCH cycle (Fig. 4b).

Finally, to show that *in vitro* self-regeneration can be performed in cell-sized compartments, we attempted to miniaturize and spatially confine these interactions in microfluidic-generated water-in-oil droplets (Fig. S16). To this end, we produced a binary emulsion of ΔEcm-MGLN, and created two droplet populations containing, or lacking DNA. Only in presence of DNA, droplets were able to perform CO_2_ fixation by autonomously producing one of the key enzymes (Ecm) necessary for their own metabolism, demonstrating that the systems established here, were also functional on a cellular scale.

Summarizing, in this study, we successfully constructed a metabolically and genetically linked network outside of a living context. We assembled this network from the bottom-up using 53 enzymes from all the domains of life, up to three DNA encoded instructions, ribosomes, tRNAs, and dozens of coenzymes, cofactors, and small molecules. Our system couples metabolism with protein production, simultaneously operates these functions *in vitro* and shows emergent properties that resemble the functions of living systems, including (self-)integration, autocatalytic feed-forward loops and (partial) self-regeneration.

While we successfully demonstrate the generation of one central building block, glycine, directly from inorganic carbon (CO_2_), a number of challenges still remain, when transitioning from such a “partial” self-regeneration towards the assembly of a system that uses CO_2_ as solely carbon source for all organic molecules. Such efforts will likely profit from recent advancements and new concepts in cell-free biology, including the development of anaplerotic reaction sequences for replenishing metabolic intermediates (*10*), improving TX-TL synthesis rates, and genetic regulation (*7*). Integrating such concepts into our platform will open new avenues for biomimetic systems that are capable of decision-making, self-repair, and eventually also self-powering from light (*17*), or even electricity (*18*) in the future.

## Acknowledgments

We thank D.G. Marchal, P.D. Gerlinger, A. Sánchez-Pascuala, D. Schindler, M. Tinzl, and H.Y. Yang for fruitful discussions. This work was supported by the Max Planck Society. S.G is grateful to the European Molecular Biology Organization (EMBO) postdoctoral fellowship (S.G. ALTF 162-2022). N.B. conducted his research within the Max Planck School Matter to Life supported by the German Federal Ministry of Education and Research (BMBF) in collaboration with the Max Planck Society.

## Funding

Open access funding was provided by the Max Planck Society.

## Author contributions

T.J.E., and S.G. conceived the work. S.G. designed the experiments; S.G., N.B., and T.J.E discussed the results. S.G., and N.B. performed cloning, enzyme activity assays, cell-free TX-TL, CETCH assays, MGLN assays, and analyzed the data. C.D., and N.B. synthesized CoA-Thioesters. S.G., C.D., and N.B. performed protein production and purification. N.P. performed mass spectrometry for amino acid analysis; N.P., N.B., and S.G. analyzed the data. T.G. performed mass spectrometry for proteomics; T.G., and S.G. analyzed the data. M.B., N.B., and S.G. performed microfluidics experiments.

T.J.E. supervised the work. S.G., and T.J.E. wrote the manuscript with contributions from all authors.

## Competing interests

The authors declare no competing interests.

## Data availability

Data supporting the findings of this study are available within the article and its Supplementary Information, or from the corresponding authors upon reasonable request.

## Materials and Methods

### Chemicals and reagents

Synthesized genes were supplied by Twist Bioscience (gBlocks), or GenScript (vectors). Primers were purchased from Sigma-Aldrich. Phusion high-fidelity DNA polymerase, Phusion HF buffer, dNTPs mix, DMSO, nuclease-free water, NTPs, and BenchMark^TM^ fluorescent protein standard were supplied by Thermo Fisher Scientific. L-glutamic dehydrogenase from bovine liver, creatine phosphokinase (CK) from rabbit muscle, carbonic anhydrase (CA) from bovine erythrocytes, protector RNase inhibitor, SigmaFAST^TM^ protease inhibitor cocktail EDTA-free, imidazole, HEPES, Sodium chloride, Magnesium acetate, Potassium glutamate, Magnesium glutamate, hemi-Magnesium glutamate, Ammonium formate, Sodium formate, Sodium bicarbonate, DTT, TCEP, creatine phosphate, folinic acid, spermidine, PLP, coenzyme B_12_, coenzyme A trilithium salt, FAD, proteinogenic amino acids,^13^C2-glycine, ^15^N-L-glutamic acid, ^13^C-Sodium bicarbonate, Iron (II) sulfate, ferric citrate, Sodium fumarate, iodoacetamide, and ReadyBlue^TM^ protein stain were purchased from Sigma-Aldrich. PURE*frex*^®^ 1.0 Solutions II (enzymes), and III (ribosomes) were supplied by Hölzel Diagnostika Handels GmbH. L-malate dehydrogenase from pig heart, and tRNAs were purchased from Roche. FluoroTect^TM^ GreenLys tRNA was provided by Promega. Glyoxylic acid was purchased from Acros organics. NADH was supplied by AppliChem GmbH. ATP, and NADPH were provided by Avantor. Sodium bicarbonate, Potassium hydroxide, Magnesium chloride, and Potassium dihydrogen phosphate were purchased from Carl ROTH GmbH. Trypsin was supplied by Serva. Acetonitrile was provided by Honeywell. QIAquick PCR purification kit was supplied by Qiagen. NucleoSpin^®^ gel, PCR clean-up, and plasmid kits were purchased from Macherey-Nagel. Precision Plus Protein^TM^ protein standard was provided by Biorad. Photoresist SU8 2015 was supplied by Kayaku Advanced Materials. PDMS SYLGARD^TM^ 184 Silicone 184 was provided by Dow. Novec^TM^ 7500 oil was purchased from 3M. Surfactant Krytox^TM^ 157 FSH was provided by miller-stephenson. All chemicals were used without any further purification. Enzymes and reagents which were indicated for storage in ultra-low temperature freezers (including PURE*frex*®) were stored at -70 °C to reduce energy consumption.

### CoA-Thioester synthesis

Propionyl-CoA and crotonyl-CoA were synthesized from their respective anhydrides according to reference (1). Ethylmalonyl-CoA was enzymatically produced from crotonyl-CoA using crotonyl-CoA carboxylase/reductase, according to reference (2). CoA-esters were purified by using HPLC 1260 Infinity (Agilent), equipped with Gemini 10 μm NX-C18 110 Å Column (Phenomenex), as described in reference (2). The concentration of CoA-esters was measured by absorbance at 260 nm with Nanodrop One^c^ (Thermo Fisher Scientific). (ε = 22.4 mM^-1^ cm^-1^, and ε = 16.4 mM^-1^ cm^-1^ for unsaturated, and saturated CoA-thioesters respectively).

### Plasmid construction

*Arabidopsis thaliana GGAT1* gene was codon optimized by Twist Bioscience for protein production in *E. coli*. The gBlock encoding GGT1 (see Table S7) was PCR amplified (using the forward primer 3 and reverse primer 4 reported in Table S6). The PCR product was gel extracted, and used to perform Gibson assembly with pET16b vector, previously amplified by PCR (with forward primer 5 and reverse primer 6 reported in Table S6). Chemically competent cells *E. coli* One Shot TOP10 (Invitrogen) were transformed with the resulting vector, and grown at 37 °C overnight on a LB agar selection plate containing Ampicillin 100 μg ml^−1^. A single colony was used to inoculate a starter culture in selective LB medium. The final construct (His-tag *GGAT1*) was extracted, and verified by sequencing (Microsynth). Other plasmids were previously constructed (Table S1).

### Protein overproduction and purification

Unless detailed in the followings, enzymes in Table S1 were overproduced and purified as described below. Chemically competent *E. coli* BL21(DE3) (ThermoScientific) cells were transformed with the plasmid encoding the respective enzyme. For Hbs production, *E. coli* BL21(DE3) (ThermoScientific) were transformed with an additional plasmid co-expressing GroEL and GroES chaperones. For Mco and Pco chemically competent *E. coli* BL21(DE3) Rosetta (Novagen) cells were used. Cells were grown at 37 °C overnight on LB agar selection plates containing Ampicillin 100 μg ml^−1^, or Streptomycin 50 μg ml^−1^ (Scr), or Chloramphenicol 34 μg ml^−1^ (KatE), or Ampicillin 100 μg ml^−1^ and Spectinomycin 50 μg ml^−1^ (Hbs). A single colony was used to inoculate a starter culture in selective LB medium. The starter culture was grown at 30 °C overnight in a shaking incubator, and used to inoculate an expression culture in selective terrific broth medium at 1:200 dilution on the next day. The expression culture was grown at 37 °C in a shaking incubator until an OD_600_ of 1 was reached. After lowering the temperature to 21 °C, the expression culture was induced with 0.5 mM isopropyl-β-d-thiogalactoside (IPTG), and grown at 21 °C in a shaking incubator overnight. For producing Hbd, the expression culture was grown into sterile Schott bottles for protein production under microaerobic conditions until an OD_600_ of 4 was reached. At the induction, cultures were further supplemented with 100 μM Fe(II)SO4, 100 μM Fe(III)citrate, and 20 mM fumarate. Cells were collected at 4,000*g* for 10 min at 10 °C, cell pellets were snap-frozen in liquid nitrogen, and stored at -20 °C until enzyme purification. Cell pellets were resuspended in twice their volume in buffer A (500 mM NaCl, 50 mM HEPES pH 7.8, glycerol 10% v/v, and one tablet of SigmaFAST protease inhibitor cocktail per liter). The cell suspension was sonicated by using Sonopuls ultrasonic homogenizer (Bandelin), equipped with KE76 probe, at 50% amplitude for lysing the cells, and centrifuged at 100,000*g* for 45 min at 4 °C. The filtered supernatant (0.20 μm filter, Sarstedt) was loaded onto 4 ml Protino Ni-NTA Agarose, in Protino gravity columns (Macherey-Nagel), previously equilibrated with 2 column volumes of buffer A. The column was washed with 2 column volumes of washing buffer (500 mM NaCl, 50 mM imidazole, 50 mM HEPES pH 7.8, glycerol 10% v/v, and one tablet of SigmaFAST protease inhibitor cocktail per liter), and the His-tagged proteins were eluted with 3 ml of buffer B (500 mM NaCl, 500 mM imidazole, 50 mM HEPES pH 7.8, glycerol 10% v/v, and one tablet of SigmaFAST protease inhibitor cocktail per liter). The eluate was concentrated with 3 kDa spin filters (Millipore), and desalted with PD-10 columns (Cytiva) and buffer C (200 mM NaCl, 50 mM HEPES pH 7.8, glycerol 10% v/v, and one tablet of SigmaFAST protease inhibitor cocktail per liter). Hbs, Pco, and Mcl1 enzymes were desalted on a size-exclusion column HiLoad 16/600 Superdex 200 pg (GE Healthcare) using buffer C. The eluted fractions were pooled, concentrated, and the purity of the purified enzymes was assessed by SDS–PAGE on 4-20% MP TGX gels (biorad), with GelStick imager (Intas), see Fig. S7. Pco and Mco enzymes were supplemented with FAD at a concentration equivalent to the protein concentration. Hbs was added with 2 mM MgCl_2_. Mcm and Ecm were supplemented with coenzyme B_12_ at a final concentration of 2 mM. Purified enzymes were added with glycerol to a final concentration of 20% v/v, snap-frozen in liquid nitrogen, and stored at −80 °C.

### Enzyme activity assays

For all enzyme assays the oxidation of NADH was detected at 340 nm on a Cary 60 UV-Vis spectrophotometer (Agilent) in quartz cuvettes with a path length of 1 cm (Hellma Optik GmbH). The enzyme activity assay for BhcA with glyoxylate and L-aspartate as substrates was performed at 30 °C in a volume of 300 μl. The reaction mixture contained 100 mM monopotassium phosphate buffer pH 7.5, 0.1 mM PLP, 0.2 mM NADH, 53 nM BhcA, and 500 nM malate dehydrogenase as a coupling enzyme to convert oxaloacetate into malate. Kinetics for glyoxylate were measured with 20 mM L-aspartate, and different concentrations of glyoxylate (5 mM, 2.5 mM, 1.25 mM, 0.63 mM, 0.31 mM, 0.16 mM). To perform the enzyme activity assay with glyoxylate and L-glutamate as substrates, the reaction mixture contained 60 mM monopotassium phosphate buffer pH 7.5, 0.1 mM PLP, 0.2 mM NADH, 53 nM BhcA, and 700 nM glutamic dehydrogenase as a coupling enzyme to convert 2-oxoglutarate into glutamate. Kinetics for glyoxylate were measured with 100 mM Potassium glutamate, and different concentrations of glyoxylate (0.5 mM, 0.25 mM, 0.13 mM, 0.06 mM, 0.03 mM, 0.015 mM). To perform the enzyme activity assay with glyoxylate and L-glutamate as substrates in the presence of PURE TX-TL buffers and salts, the reaction mixture contained 50 mM HEPES buffer pH 7.6, 0.1 mM PLP, 0.2 mM NADH, 11.8 mM Magnesium acetate, 0.3 mM each of the 19 proteinogenic AAs (-glycine), 53 nM BhcA, and 700 nM glutamic dehydrogenase. Kinetics for glyoxylate were measured with 100 mM Potassium glutamate, and different concentrations of glyoxylate (0.5 mM, 0.25 mM, 0.13 mM, 0.06 mM, 0.03 mM, 0.015 mM). The enzyme activity assay for GGT1 with glyoxylate and L-glutamate as substrates was performed at 30 °C in a volume of 300 μl. The reaction mixture contained 60 mM monopotassium phosphate buffer pH 7.5, 0.1 mM PLP, 0.2 mM NADH, 7.6 nM GGT1, and 700 nM glutamic dehydrogenase as a coupling enzyme to convert 2-oxoglutarate into glutamate. Kinetics for glyoxylate were measured with 20 mM L-glutamate, and different concentrations of glyoxylate (5 mM, 2.5 mM, 1.25 mM, 0.63 mM, 0.31 mM, 0.16 mM). To perform the enzyme activity assay with glyoxylate and L-glutamate as substrates in the presence of PURE TX-TL buffers and salts, the reaction mixture contained 50 mM HEPES buffer pH 7.6, 0.1 mM PLP, 0.2 mM NADH, 11.8 mM Magnesium acetate, 0.3 mM 19 proteinogenic AAs (-glycine), 7.6 nM GGT1, and 700 nM glutamic dehydrogenase as a coupling enzyme to convert 2-oxoglutarate into glutamate. Kinetics for glyoxylate were measured with 100 mM Potassium glutamate, and different concentrations of glyoxylate (5 mM, 2.5 mM, 1.25 mM, 0.63 mM, 0.31 mM, 0.16 mM). The Michaelis–Menten kinetics of the transamination reactions in this study are reported in Fig. S1, and the calculated kinetic parameters are provided in Table S2.

### Mass spectrometry enzyme assays

For all enzyme assays the concentration of ^15^N-glycine and glycine were quantified at several time points by HPLC-MS/MS while monitoring the concentration of the other amino acids, except for glutamate (amino-group donor present at high concentration), and cysteine (prone to oxidation). For alanine, asparagine, aspartic acid, phenylalanine, serine, tryptophan, and tyrosine ^15^N-residues were also quantified.

#### Reaction assembly

The enzyme assay for BhcA with glyoxylate and L-glutamate as substrates was performed at 30 °C for 15 min, shaking (500 rpm), in Thermomixer pro (CellMedia), in a volume of 600 μl. The reaction mixture contained 50 mM HEPES buffer pH 7.6, 0.1 mM PLP, 11.8 mM Magnesium acetate, 0.3 mM 19 proteinogenic AAs (-glycine), 0.3 mM glyoxylate, 100 mM ^15^N-Potassium glutamate, and 53 nM BhcA. To perform the enzyme assay for GGT1 with glyoxylate and L-glutamate as substrates, 7.6 nM, or 76 nM GGT1 were used. Negative controls without enzymes (BhcA, or GGT1), or substrate (glyoxylate) were measured. Additional enzyme assays for BhcA, or GGT1 with glyoxylate and L-glutamate as substrates were performed at 30 °C for 4 h, shaking (500 rpm), in Thermomixer pro (CellMedia), in a volume of 1200 μl.

#### Sampling

Dependent on the assay, samples were taken at different time points representing different reaction times. At the sampling point, assays were stopped by filtration at 14,000*g* and 4 °C with 10 kDa spin filters (Millipore).

#### Sample preparation for HPLC-MS/MS quantification

After filtration samples were diluted 1/9 v/v sample:dilution solution (50 mM HEPES buffer pH 7.6, 0.1 mM PLP, 11.8 mM Magnesium acetate, 100 mM Potassium glutamate) prior to mass spectrometry. 20 amino acids standards for calibration curves were prepared at 1 μM, 2.5 μM, 5 μM, 10 μM, 25 μM, 50 μM, and 100 μM in calibration solution (50 mM HEPES buffer pH 7.6, 0.1 mM PLP, 11.8 mM Magnesium acetate, 0.3 mM glyoxylate, 100 mM ^15^N-Potassium glutamate) to account for matrix effects.

#### HPLC-MS/MS quantification

Quantitative determination of amino acids was performed by using HPLC-MS/MS. The chromatographic separation was performed on Infinity II 1290 HPLC system (Agilent) using a ZicHILIC SeQuant column (150 × 2.1 mm, 3.5 μm particle size, 100 Å pore size) connected to a ZicHILIC guard column (20 × 2.1 mm, 5 μm particle size) (Merck KgAA), with a constant flow rate of 0.3 ml min^−1^, with mobile phase A being 0.1% formic acid in 99/1 v/v water:acetonitrile and phase B being 0.1% formic acid in 1/99 v/v water:acetonitrile at 25 °C. The injection volume was 1 μl. The mobile phase profile consisted of the following steps and linear gradients: 0 – 8 min from 80 to 60% B; 8 – 10 min from 60 to 10% B; 10 – 12 min constant at 10% B; 12 – 12.1 min from 10 to 80% B; 12.1 to 14 min constant at 80% B. An Agilent Triple Quad 6495 ion funnel mass spectrometer was used in positive mode with an electrospray ionization source and the following conditions: ESI spray voltage 2000 V, nozzle voltage 1000 V, sheath gas 250 °C at 12 l min^−1^, nebulizer pressure 60 psig, and drying gas 100 °C at 11 l min^−1^. Compounds were detected using dynamic multi reaction monitoring, and identified based on their mass transition and retention time compared to standards. Chromatograms were integrated by using MassHunter software (Agilent, Santa Clara, CA, USA), and the Agile2 integrator. Labeling distributions were determined using the same instrumentation and chromatographical method based on peak areas. Absolute concentrations of labeled and unlabeled amino acids were calculated based on an external calibration curve of the respective unlabeled amino acid prepared in sample matrix. The mass transitions, collision energies, fragmentor and accelerator voltages are reported in Table S3. Shown are bar-plots of the statistical mean of the results of triplicates (Figs. S2-S6); error bars represent the standard deviation of the same data.

### Cell-Free TX-TL

#### Energy solution preparation

SolutionA(-salts - tRNAs - AAs), salts solution, and tRNAs solution were prepared as described in reference (3), and mixed 3.4/1.6/2.5 v/v/v solutionA(-salts - tRNAs - AAs):salts solution:tRNAs solution to reach 20 mM creatine phosphate, 0.02 mM folinic acid, 2 mM spermidine, 1 mM DTT, 2 mM ATP, 2 mM GTP, 1 mM CTP, 1 mM UTP, 50 mM HEPES buffer pH 7.6, 11.8 mM Magnesium acetate, 100 mM Potassium glutamate, and 56 A_260_ ml^−1^ tRNAs in TX-TL reactions.

#### Control of TX-TL with glycine

A Δgly-PURE TX-TL system was assembled by mixing 3.4 μl solutionA(-Salts tRNAs - AAs), 1.6 μl salts solution, 2.5 μl tRNAs solution,1.25 μl PURE*frex* Solutions II (enzymes), 1.25 μl PURE*frex* Solutions III (ribosomes), 0.5 μl RNAse inhibitor, 75 ng *EGFP* template (previously PCR-amplified using *EGFP* gBlock, forward primer 1 and reverse primer 2 reported in Tables S7 and S6 respectively), and 2.5 μl 19 proteinogenic AAs (-glycine), 3 mM each. Reactions were mixed in ice, and nuclease-free water was added to bring the reaction volume to 25 μl. Glycine was added at a final concentration of 300 μM, or at a final concentration ranging from 10 μM to 300 μM (10 μM, 19 μM, 38 μM, 75 μM, 150 μM, and 300 μM) in TX-TL reactions. Negative controls without glycine were performed. Reactions were gently mixed, transferred into a Nunc^TM^ 384-well black, transparent bottom, plate (Thermo Fisher Scientific), sealed with SealPlate film (Sigma-Aldrich), centrifuged at 2,000*g* and 4 °C, and incubated at 37 °C in Infinite M Plex microplate reader (Tecan) for 6 h. The plate reader was operated in fluorescent mode, bottom reading, λ_exc_ = 488 nm, λ_em_ = 515 nm, 1 min interval read, 20s orbital shaking, and gain = 100. Shown are plots of the statistical mean of the results of triplicates at any acquisition time; error bars represent the standard deviation of the same data.

#### Control of TX-TL with glyoxylate and aminotransferase enzyme

Δgly-PURE TX-TL system was supplemented with 0.1 mM PLP, 7.6 nM GGT1, and glyoxylate at a final concentration of 300 μM, or at a final concentration ranging from 10 μM to 300 μM (10 μM, 19 μM, 38 μM, 75 μM, 150 μM, and 300 μM) in 25 μl TX-TL reactions. Negative controls without enzyme (GGT1) and substrate (glyoxylate), or without GGT1, or without glyoxylate were measured. A reference control with glycine at a final concentration of 300 μM in TX-TL reaction was also performed. The microplate reader was operated with gain = 90.

#### Control of TX-TL with glycine in presence of CETCH components

Δgly-PURE TX-TL system was supplemented with CETCH components (15 mM HEPES buffer pH 7.8, 2.5 mM MgCl_2_, 1 mM CP, 2 mM Sodium bicarbonate, 8 mM Sodium formate, 0.1 mM CoA, 2 mM ATP, 0.75 mM NADPH, 0.5 μM Pco, 0.4 μM Ccr, 0.3 μM Epi, 0.3 μM Mcm, 1.4 μM Scr, 0.9 μM Ssr, 0.1 μM Hbs, 0.3 μM Hbd, 0.6 μM Ecm, 9.3 μM Mco, 0.1 μM Mch, 2.9 μM Mcl1, 0.3 μM KatE, 2.6 μM Fdh, 0.5 μM CK, 6 nM CA) or (30 mM HEPES buffer pH 7.8, 5 mM MgCl_2_, 2 mM CP, 4 mM Sodium bicarbonate, 16 mM Sodium formate, 0.2 mM CoA, 4 mM ATP, 1.5 mM NADPH, 0.9 μM Pco, 0.9 μM Ccr, 0.6 μM Epi, 0.6 μM Mcm, 2.8 μM Scr, 1.8 μM Ssr, 0.2 μM Hbs, 0.6 μM Hbd, 1.2 μM Ecm, 18.6 μM Mco, 0.2 μM Mch, 5.9 μM Mcl1, 0.7 μM KatE, 5.2 μM Fdh, 1.1 μM CK, 13 nM CA), 20 μM or 40 μM PLP, 20 nM or 40 nM GGT1, and glycine at a final concentration of 300 μM in 25 μl TX-TL reactions. Negative controls without glycine, or without glycine and DNA template were performed. Reactions were incubated at 37 °C into the microplate reader for 16 h, with gain = 90.

#### Cell-free TX-TL of GGT1, Ecm, and Epi enzymes

A PURE TX-TL system was assembled by mixing 3.4 μl solutionA(-Salts tRNAs - AAs), 1.6 μl salts solution, 2.5 μl tRNAs solution,1.25 μl PURE*frex* Solutions II (enzymes), 1.25 μl PURE*frex* Solutions III (ribosomes), 0.5 μl RNAse inhibitor, 2.5 μl 20 proteinogenic AAs (3 mM each), 1 μl of FluoroTect GreenLys tRNA, and 75 ng of *GGAT1*, or *Ecm*, or *Epi* templates (previously PCR-amplified using pUC57_*GGAT1*, pUC57_*Ecm* , or pUC57_*Epi* vectors respectively, and forward primer 1 and reverse primer 2 reported in Tables S7 and S6). Nuclease-free water was added to bring the reaction volume to 25 μl. Reactions were gently mixed, incubated at 37 °C in Thermomixer pro (CellMedia) for 4h, analyzed by SDS–PAGE on 4-20% MP TGX gels (Biorad), and imaged with ChemiDoc system (Biorad), see Fig. S8. The activity of GGT1 was assayed by co-expression of *GGAT1* and *EGFP* DNA templates in Δgly-PURE TX-TL system. The assay was run in Infinite M Plex microplate reader (Tecan), and Δgly-PURE TX-TL system was supplemented with 0.1 mM PLP, 0.3 mM glyoxylate, 10 μM glycine, and 50 ng of each DNA template. Negative controls without glyoxylate, or without glyoxylate and glycine were performed. An additional control with 0.3 mM glyoxylate, 10 μM glycine and without *GGAT1* template was included. The activity of Ecm was tested by co-expression of *Ecm* and *EGFP* DNA templates in Δgly-PURE TX-TL system. The assay was run in the microplate reader, and Δgly-PURE TX-TL system was supplemented with 15 mM HEPES buffer pH 7.8, 2.5 mM MgCl_2_, 1 mM CP, 2 mM Sodium bicarbonate, 8 mM Sodium formate, 0.1 mM CoA, 2 mM ATP, 0.75 mM NADPH, 0.1 mM PLP, 0.3 μM Epi, 9.3 μM Mco, 0.1 μM Mch, 2.9 μM Mcl1, 0.3 μM KatE, 20 nM GGT1, 100 μM ethylmalonyl-CoA, 10 μM glycine, and 50 ng of each DNA template. Negative controls without ethylmalonyl-CoA, or without ethylmalonyl-CoA and glycine were performed. The activity of Epi was tested by co-expression of *Epi* and *EGFP* DNA templates in Δgly-PURE TX-TL system. The assay was run in the microplate reader, and Δgly-PURE TX-TL system was supplemented with 15 mM HEPES buffer pH 7.8, 2.5 mM MgCl_2_, 1 mM CP, 2 mM Sodium bicarbonate, 8 mM Sodium formate, 0.1 mM CoA, 2 mM ATP, 0.75 mM NADPH, 0.1 mM PLP, 0.6 μM Ecm, 9.3 μM Mco, 0.1 μM Mch, 2.9 μM Mcl1, 0.3 μM KatE, 20 nM GGT1, 100 μM ethylmalonyl-CoA, 10 μM glycine, and 25 ng of *Epi* and 50 ng of *EGFP* templates respectively. Negative controls without ethylmalonyl-CoA, or without ethylmalonyl-CoA and glycine were performed. Reactions were incubated at 30 °C into the microplate reader for 6 h, with gain = 100, see Figs. S10-12.

#### Cell-free TX-TL with different Mg^2+^ ion sources

A PURE TX-TL system was assembled by mixing 3.4 μl solutionA(-Salts tRNAs - AAs), 2.5 μl tRNAs solution, 1.25 μl Potassium glutamate (2M), 1.25 μl PURE*frex* Solutions II (enzymes), 1.25 μl PURE*frex* Solutions III (ribosomes), 0.5 μl RNAse inhibitor, 2.5 μl 20 proteinogenic AAs (3 mM each), 75 ng *EGFP* template (previously PCR-amplified using *EGFP* gBlock, and forward primer 1 and reverse primer 2 reported in Tables S6 and S7), and different sources of Mg^2+^ ion (11.8 mM Magnesium acetate, or 11.8 mM hemi-Magnesium glutamate, or 11.8 mM MgCl_2_, or 5.9 mM Magnesium acetate and 5.9 mM MgCl_2_, or 2.5 mM MgCl_2_). Negative controls without any source of Mg^2+^ were also tested. Reactions were incubated at 37 °C into the microplate reader for 6 h, with gain = 90.

#### Cell-free TX-TL at low acetate concentration

To further decrease the concentration of acetate in TX-TL reactions, a buffer exchanged PURE TX-TL system was obtained by mixing 3.4 μl solutionA(-Salts tRNAs - AAs), 2.5 μl tRNAs solution, 1.25 μl Potassium glutamate (2M), 1.25 μl buffer exchanged PURE*frex* Solutions II (enzymes), 1.25 μl buffer exchanged PURE*frex* Solutions III (ribosomes), 0.5 μl RNAse inhibitor, 2.5 μl 20 proteinogenic AAs (3 mM each), 75 ng *EGFP* template, and different sources of Mg^2+^ ion (described above). Buffer exchanged PURE*frex* Solutions II (enzymes) and III (ribosomes) were obtained by buffer exchanging the enzymes and the ribosomes of PURE*frex* with buffer D (50 mM HEPES buffer pH 7.6, 100 mM Potassium glutamate, 11.8 mM hemi-Magnesium glutamate) and buffer E (50 mM HEPES buffer pH 7.6, 30 mM Potassium glutamate, 40 mM hemi-Magnesium glutamate) respectively. 3 kDa spin concentrators (Millipore) were used, and glycerol was added to buffer exchanged enzymes at final concentration of 30% v/v. Negative controls without any source of Mg^2+^ were also tested. Reactions were incubated at 37 °C into the microplate reader for 6 h, with gain = 90, see Fig. S9.

### CETCH assays

For all CETCH assays the reaction assembly is based on CETCH (75 mM HEPES buffer pH 7.8, 12.5 mM MgCl_2_, 5 mM CP, 10 mM Sodium bicarbonate, 40 mM Sodium formate, 0.5 mM CoA, 10 mM ATP, 3.75 mM NADPH, 2.3 μM Pco, 2.2 μM Ccr, 1.5 μM Epi, 1.5 μM Mcm, 7 μM Scr, 4.4 μM Ssr, 0.5 μM Hbs, 1.5 μM Hbd, 2.9 μM Ecm, 46.5 μM Mco, 0.3 μM Mch, 14.7 μM Mcl1, 1.6 μM KatE, 13.1 μM Fdh, 2.75 μM CK, 30 nM CA) recently optimized by using active learning (4). To produce glycine as output of the CETCH cycle, the assay was supplemented with PLP, GGT1, and different concentrations of propionyl-CoA.

#### Glycine production by CETCH in presence of PURE TX-TL components

Mixtures 1/4 v/v CETCH:dilution solution were assembled to achieve the final compositions reported in Table S5. All the assays were provided with 20 μM PLP, and 20 nM GGT1, and started with 200 μM propionyl-CoA. Mixtures were incubated at 30 °C for 4h, shaking (500 rpm), in Thermomixer pro (CellMedia). Assays were stopped by filtration at 14,000*g* and 4 °C with 3 kDa spin filters (Millipore). After filtration, samples were diluted, mixed 1/1 v/v sample:internal standard (5 μM ^13^C2-glycine) prior to mass spectrometry to account for matrix effects. Glycine standards were prepared at 1 μM, 2.5 μM, 5 μM, 10 μM, 25 μM, 50 μM, and 100 μM in water, and diluted 1/1 v/v calibrant:internal standard. For all assays glycine quantification was performed by HPLC-MS/MS using the method described above, with multi reaction monitoring (see Tables S3-4).

### MGLN assays

#### Control of TX-TL with CO_2_

MGLNs were assembled in 25 μl reaction volumes by supplementing a buffer exchanged Δgly-PURE TX-TL system with 11.8 hemi-Magnesium glutamate, 15 mM HEPES buffer pH 7.8, 2.5 mM MgCl_2_, 1 mM CP, 2 mM Sodium bicarbonate, 8 mM Sodium formate, 0.1 mM CoA, 2 mM ATP, 0.75 mM NADPH, 20 μM PLP, 0.5 μM Pco, 0.4 μM Ccr, 0.3 μM Epi, 0.3 μM Mcm, 1.4 μM Scr, 0.9 μM Ssr, 0.1 μM Hbs, 0.3 μM Hbd, 0.6 μM Ecm, 9.3 μM Mco, 0.1 μM Mch, 2.9 μM Mcl1, 0.3 μM KatE, 2.6 μM Fdh, 0.5 μM CK, 6 nM CA, and 20 nM GGT1. 75 ng EGFP template, and different concentrations of propionyl-CoA (200 μM, 100 μM, and 50 μM) were added. Negative controls without propionyl-CoA, or without propionyl-CoA and DNA template were performed. MGLN were incubated at 30 °C into the microplate reader for 16 h, with gain = 100.

#### Autocatalytic MGLN

A ΔGGT1-MGLN was assembled in a reaction volume of 25 μl, as described above, without adding the aminotransferase enzyme. The ΔGGT1-MGLN was supplemented with 10 μM glycine, 50 ng *GGAT1* template, 50 ng *EGFP* template, and 50 μM propionyl-CoA. Negative controls without propionyl-CoA, or without propionyl-CoA and glycine, or without DNA templates were performed. MGLNs were incubated at 30 °C into the microplate reader for 16 h, with gain = 100.

#### Self-regenerating MGLN

A ΔEcm-MGLN was assembled in a reaction volume of 25 μl, as described above, without adding the Ecm enzyme. The ΔEcm-MGLN was supplemented with 10 μM glycine, 6 μM coenzyme B_12_, 50 ng *Ecm* template, 50 ng *EGFP* template, and different concentrations of propionyl-CoA (200 μM, 100 μM, and 50 μM). Negative controls without propionyl-CoA, or without propionyl-CoA and glycine, or without DNA templates were performed. MGLN were incubated at 30° C into the microplate reader for 16 h, with gain = 100. Self-regenerating ΔEcm-MGLN has been scaled up, and performed with CO_2_ (from NaHCO_3_), or ^13^CO_2_ (from NaH^13^CO_3_); His-tag proteins have been purified by using MagneHis system (Promega) and DynaMag spin magnet (Thermo Fisher Scientific), and submitted for proteomic analysis. A Δ(Ecm,Epi)-MGLN was further assembled, without adding Ecm and Epi enzymes. The Δ(Ecm,Epi)-MGLN was supplemented with 10 μM glycine, 6 μM coenzyme B_12_, 50 ng *Ecm* template, 25 ng *Epi* template, 50 ng *EGFP* template, and 50 μM propionyl-CoA. Negative controls without propionyl-CoA, or without propionyl-CoA and glycine, or without DNA templates were performed.

#### Shotgun proteomics for analyzing ^13^C-glycine incorporation

Protein reduction was performed in presence of 5 mM TCEP for 15 min at 90 °C, followed by alkylation using 10 mM iodoacetamide at 25 °C for 30 min. Further protein cleanup and tryptic digest were performed with SP3 beads (GE Healthcare), using the SP3 protocol described in reference (5). 1 µg trypsin was used for protein digestion, carried out overnight at 30 °C. After digest, peptides were desalted by using C18 solid phase extraction cartridges (Macherey-Nagel). Cartridges were prepared by adding acetonitrile, followed by equilibration with 0.1% TFA. Peptides were loaded on equilibrated cartridges, washed with 19/1 v/v water:acetonitrile and 0.1% TFA containing buffer, and finally eluted with 1/1 v/v water:acetonitrile and 0.1% TFA. Dried peptides were reconstituted in 0.1% TFA, and analyzed by liquid-chromatography-mass spectrometry on Orbitrap Exploris 480 (Thermo Fisher Scientific) equipped with an Ultimate 3000 RSLCnano system (Thermo Fisher Scientific), and Nanospray Flex ion source (Thermo Fisher Scientific). Peptide separation was performed on a reverse phase HPLC column (75 μm x 42 cm) packed in-house with C18 resin (2.4 μm; Dr. Maisch). The following separating gradient was used: 94% solvent A (0.15% formic acid) and 6% solvent B (0.15% formic acid in acetonitrile) to 35% solvent B over 60 minutes respectively, at 300 nl min^−1^ flow rate. Peptides were ionized at 2.3 kV spray voltage, ion transfer tube temperature was set at 275 °C, 445.12003 m/z was used as internal calibrant. Data acquisition mode was set to obtain one high resolution MS scan (60,000 full width at half maximum at m/z 200) followed by MS/MS scans of the 10 and 20 most intense ions respectively. To increase the efficiency of MS/MS attempts the charged state screening modus was enabled to exclude unassigned and singly charged ions. The dynamic exclusion duration was set to 14 s; the ion accumulation time was set to 50 ms (MS) and 50 ms at 17,500 resolution (MS/MS). The automatic gain control (AGC) was set to 3 x 106 for MS survey scan and 2 x 105 for MS/MS scans. The quadrupole isolation was 1.5 m/z; collision was induced with an HCD collision energy of 27%. MS raw data were analyzed with MaxQuant (version 2.0.3.0), and fasta database containing target proteins. MaxQuant was executed in standard settings. The search criteria were set as follows: full tryptic specificity was required (cleavage after lysine or arginine residues); two missed cleavages were allowed; carbamidomethylation (C) was set as fixed modification; oxidation (M), deamidation (N,Q), and Label:1C13 (G) variable modification. Intensities of 1C13 labeled, and unlabeled peptides were analyzed (see Fig. S15).

### Microfluidics

#### Droplet production

Droplet production was carried out using the chip design from the Drop-seq protocol described in reference (6). The mold was prepared by using photolithography, with SU8 2015 photoresist, spun at 1000 rpm for 30 s, as in reference (7). PDMS was poured on the mold, allowed to cure, and peeled off; 0.75 mm holes were punched through each of the inlets. The chip was bonded onto a glass slide by oxygen plasma treatment for 30 s. The chip was treated with ArmorAll Shield glass protector (Energizer) to produce a hydrophobic surface. Samples, or negative controls were injected in one aqueous inlet at 1 mL h^-1^. The remaining aqueous inlet was plugged with a paperclip. Sample contained the ΔEcm-MGLN, while the negative control lacked DNA templates. 1 mM Cascade Blue^TM^ hydrazide trilithium salt dye (Thermo Fisher Scientific) was added for barcoding to the negative control reaction. In the oil inlet, Novec^TM^ 7500 oil with 2.2 % v/v Krytox^TM^ 157 FSH was injected at 1.75 ml h^-1^ for stable droplet production.

#### Imaging

Droplets were incubated at 30 °C in a Thermomixer pro (CellMedia), and imaged with the 4x objective every 2 hours on a glass slide using brightfield illumination and fluorescence channels for DAPI and GFP on a Ti2-E microscope (Nikon). Image processing was performed by using Fiji/ImageJ.

---

**Fig. S1.**
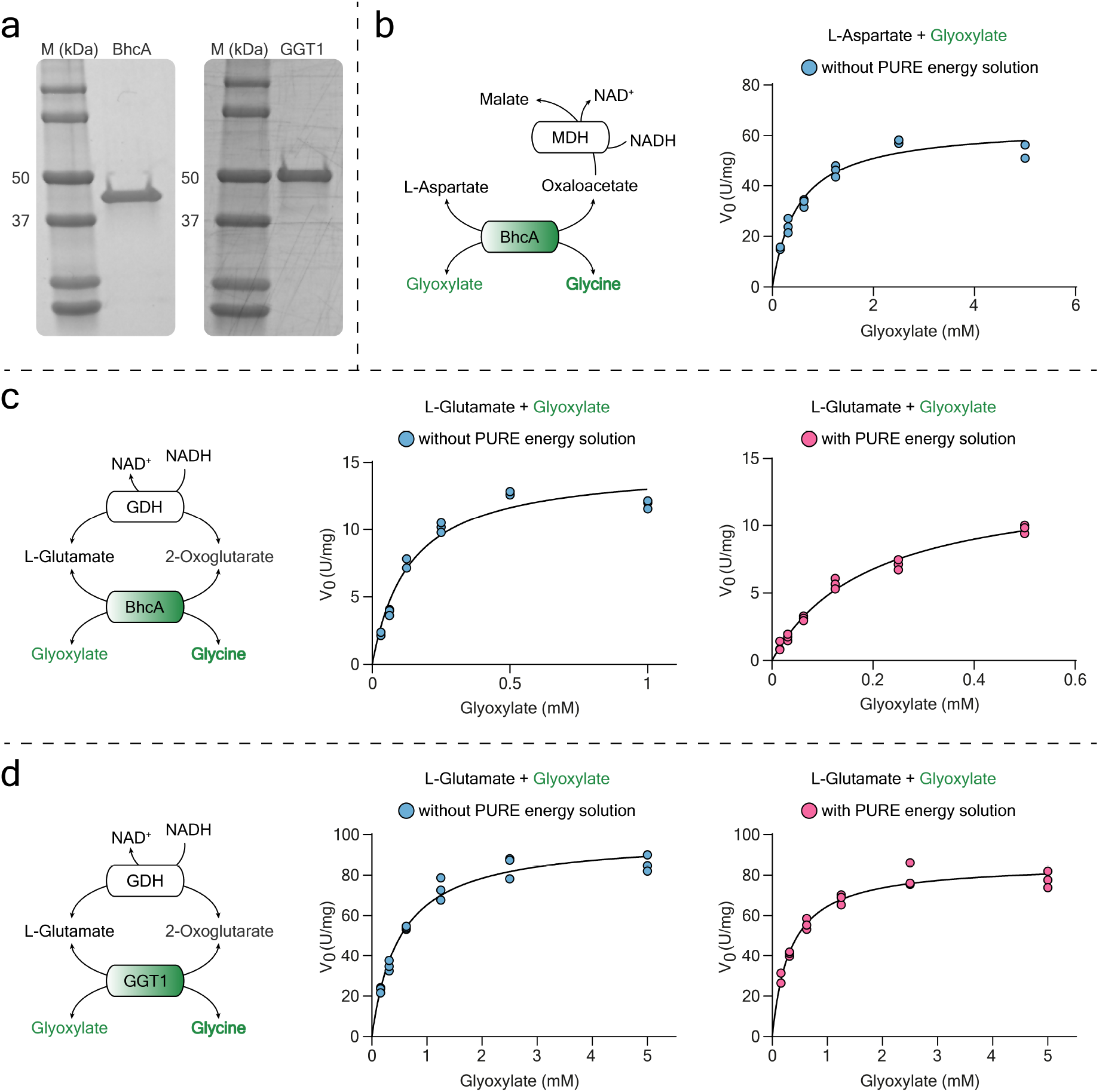
Michaelis–Menten kinetics of the transamination reactions in this study. (**a**) SDS-PAGE of purified aspartate-glyoxylate aminotransferase (BhcA, 45.0 kDa) and glutamate-glyoxylate aminotransferase (GGT1, 55.8 kDa) on 4-20% gels. (**b**) Michaelis-Menten kinetics for BhcA with aspartate. (**c**) Michaelis-Menten kinetics for BhcA with glutamate. (**d**) Michaelis-Menten kinetics for GGT1 with glutamate. **(b**-**d)** Data obtained from 3 independent experiments, at different glyoxylate concentrations, are summarized in Table S2.

**Fig. S2.**
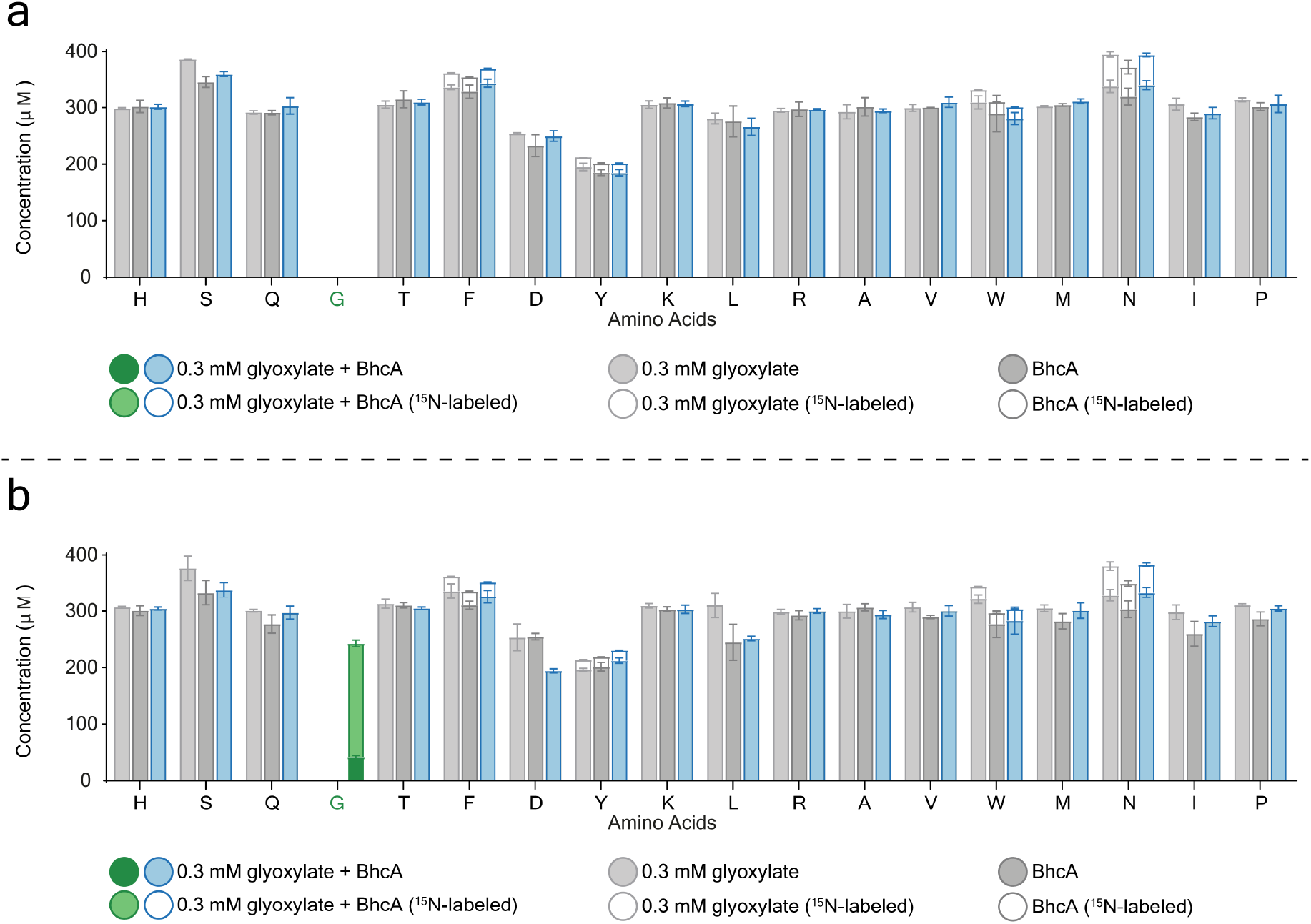
Amino acid analysis. Bar graphs showing the result of the amino acid analysis performed using mass spectrometry of the transamination reaction with BhcA aminotransferase, 0.3 mM glyoxylate and 100 mM ^15^N-glutamate substrates, in presence of the 19 proteinogenic amino acids (glycine omitted). The assay was stopped at time15 min, and the AAs were quantified at time 0 min and 15 min as shown in (**a**) and (**b**) respectively. Colored bars represent the assay performed in presence of both enzyme and substrate. Glycine is highlighted in green. Light, and dark grey bars represent the assays performed without BhcA, or glyoxylate respectively.

**Fig. S3.**
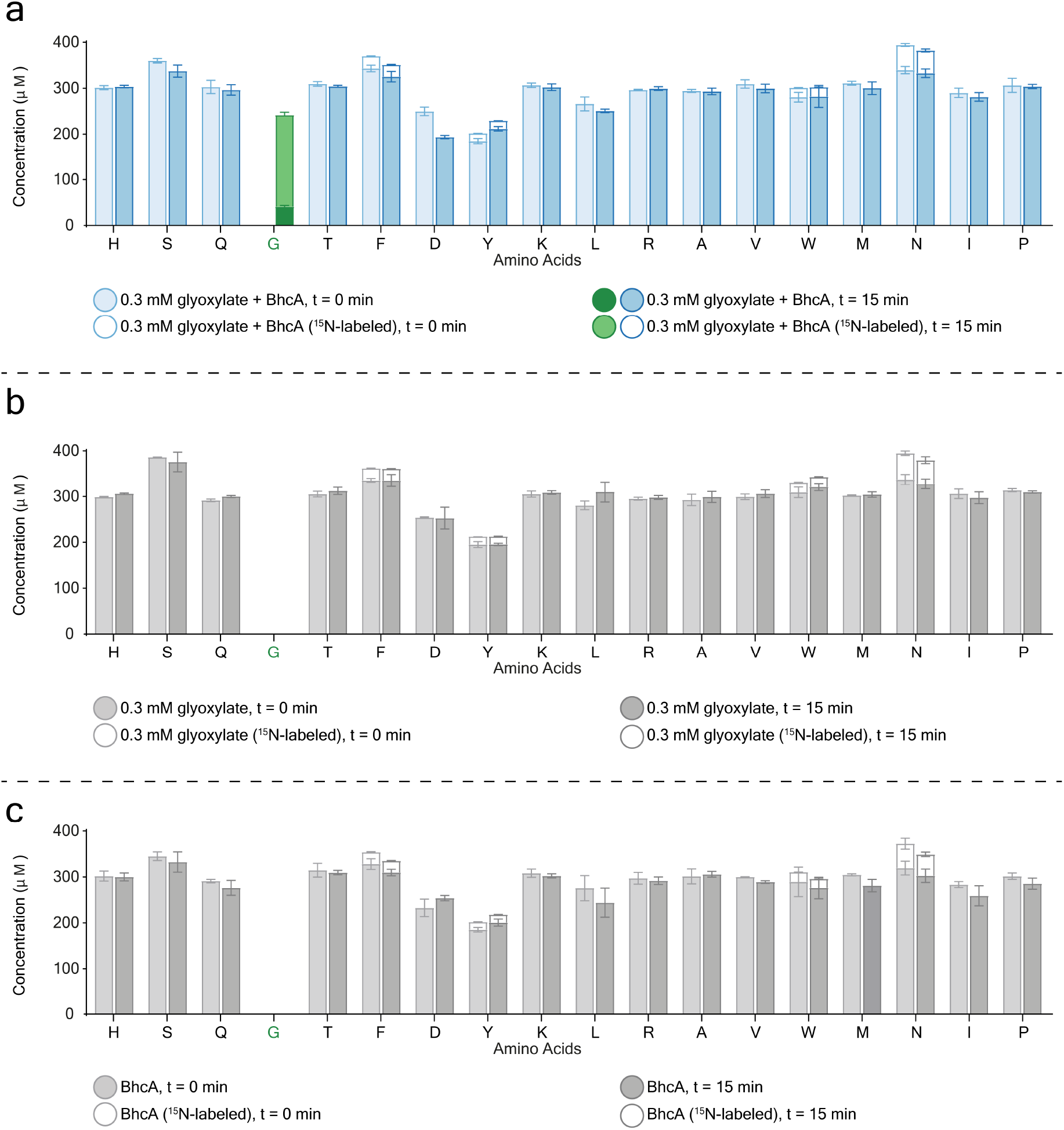
Amino acid analysis. Bar graphs showing the result of the amino acid analysis performed using mass spectrometry of the transamination reaction with BhcA aminotransferase, 0.3 mM glyoxylate and 100 mM ^15^N-glutamate substrates, in presence of the 19 proteinogenic amino acids (glycine omitted). The assay was run for 15 min, and plotted is the comparison between the concentration of each AA at time 0 min, and at time 15 min. In (**a**) the transamination reaction is performed in presence of both enzyme and substrate. In (**b**) the assay is run without BhcA. In (**c**) the reaction is performed without glyoxylate.

**Fig. S4.**
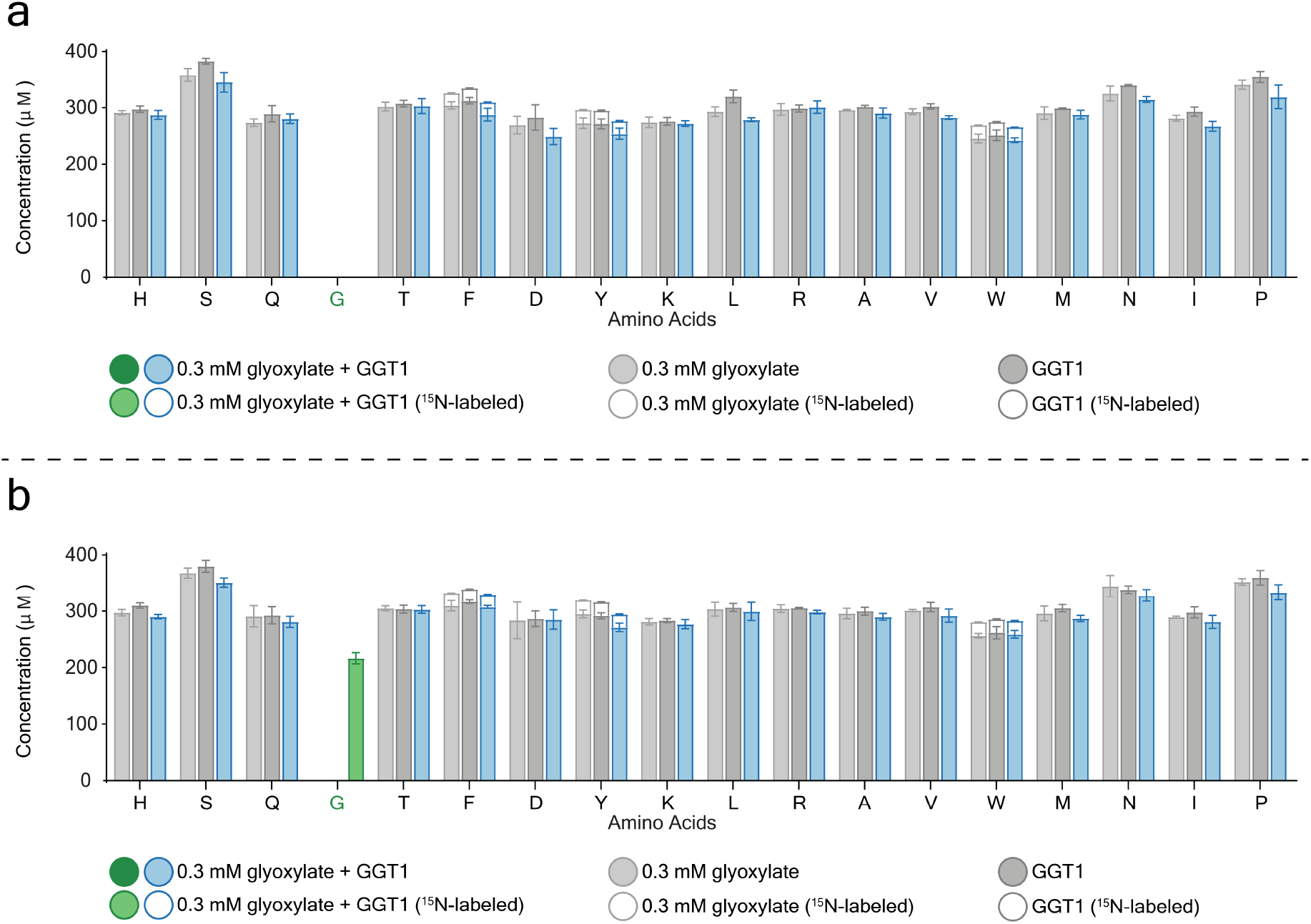
Amino acid analysis. Bar graphs showing the result of the amino acid analysis performed using mass spectrometry of the transamination reaction with GGT1 aminotransferase, 0.3 mM glyoxylate and 100 mM ^15^N-glutamate substrates, in presence of the 19 proteinogenic amino acids (glycine omitted). The assay was stopped at time15 min, and the AAs were quantified at time 0 min and 15 min as shown in (**a**) and (**b**) respectively. Colored bars represent the assay performed in presence of both enzyme and substrate. Glycine is highlighted in green. Light, and dark grey bars represent the assays performed without GGT1, or glyoxylate respectively.

**Fig. S5.**
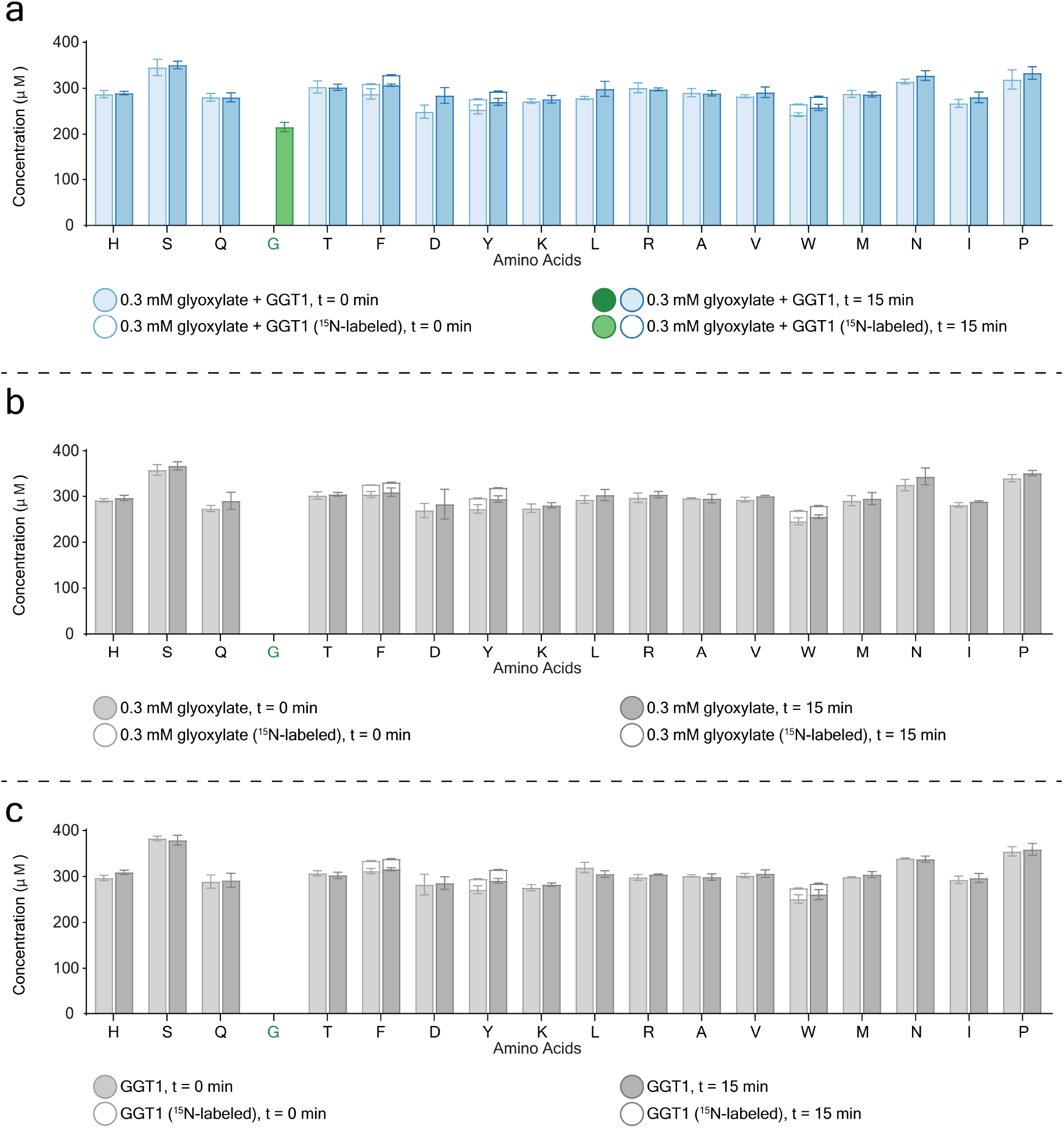
Amino acid analysis. Bar graphs showing the result of the amino acid analysis performed using mass spectrometry of the transamination reaction with GGT1 aminotransferase, 0.3 mM glyoxylate and 100 mM ^15^N-glutamate substrates, in presence of the 19 proteinogenic amino acids (glycine omitted). The assay was run for 15 min, and plotted is the comparison between the concentration of each AA at time 0 min, and at time 15 min. In (**a**) the transamination reaction is performed in presence of both enzyme and substrate. In (**b**) the assay is run without GGT1. In (**c**) the reaction is performed without glyoxylate.

**Fig. S6.**
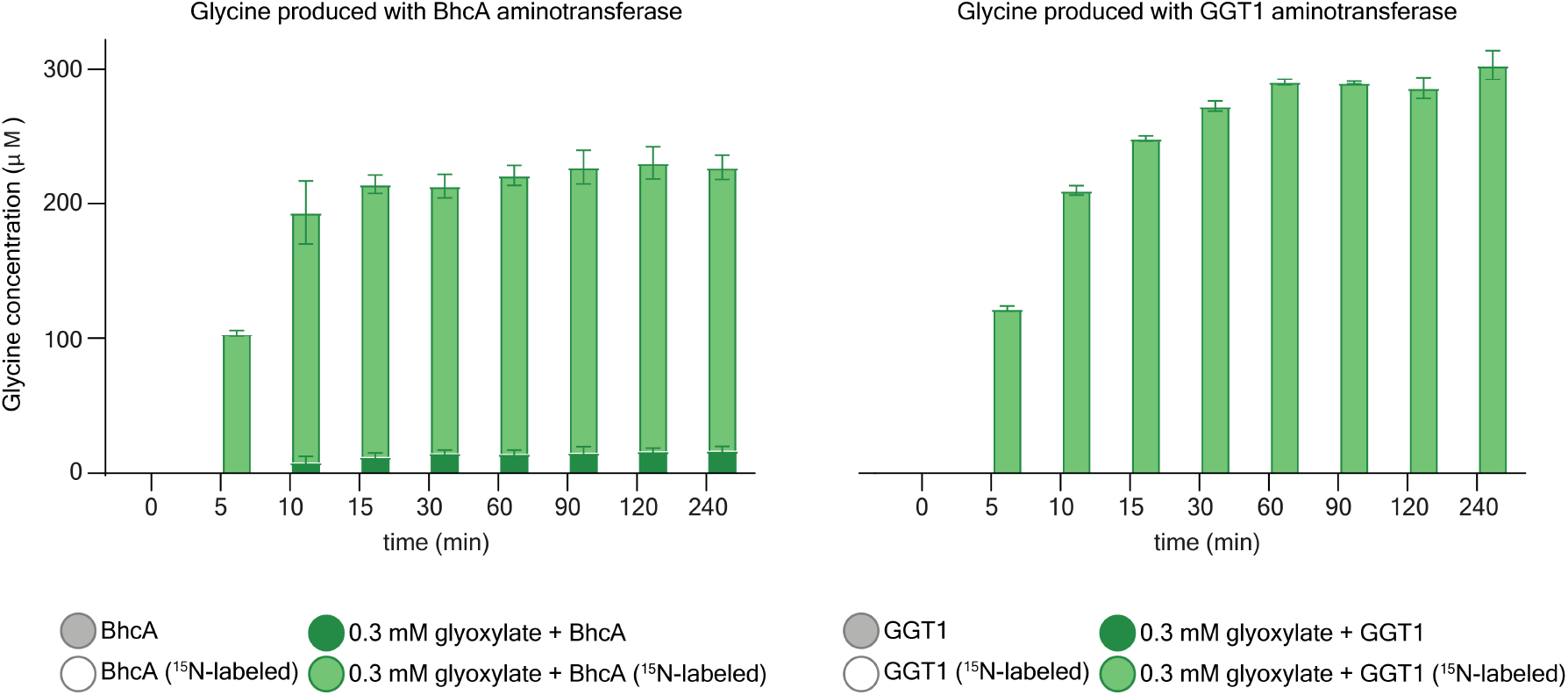
Amino acid analysis. Bar graphs showing the result of glycine quantification performed by mass spectrometry of the transamination reaction with BhcA or GGT1 aminotransferases, 0.3 mM glyoxylate and 100 mM ^15^N-glutamate substrates, in presence of the 19 proteinogenic amino acids (glycine omitted), at different time points. For each time point the concentration of glycine produced by the transamination reaction supplemented with enzyme and substrate is compared with the concentration of glycine obtained by performing the same reaction without substrate (glyoxylate).

**Fig. S7.**
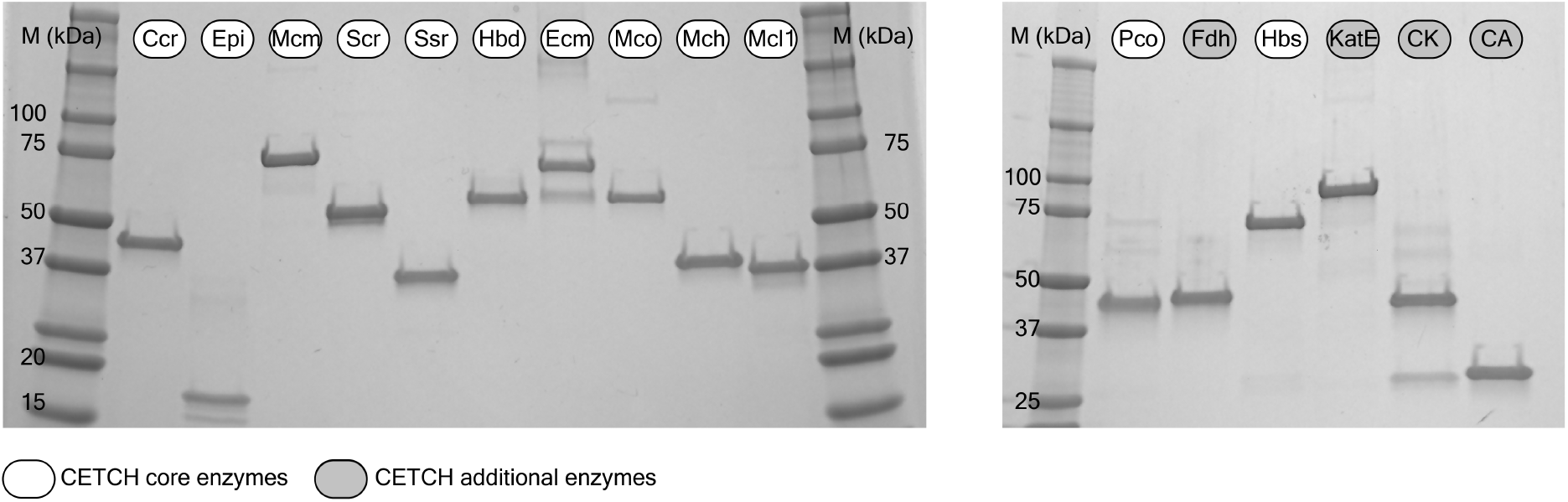
SDS PAGE of the enzymes of the CETCH cycle. The purity of all the enzymes of the CETCH cycle was assessed by SDS–PAGE on 4-20% gels. Expected monomeric molecular masses for the proteins: Ccr (49.0 kDa), Epi (16.8 kDa), Mcm (80.1 kDa), Scr (51.5 kDa), Ssr (39.2 kDa), Hbd (60.5 kDa), Ecm (73.7 kDa), Mco (62.2 kDa), Mch (39.7 kDa), Mcl1 (36.8 kDa), Pco (50.1 kDa), Fdh (46.6 kDa), Hbs (78.1 kDa), KatE (86.5 kDa), CK (43.1 kDa), CA (30.0 kDa).

**Fig. S8.**
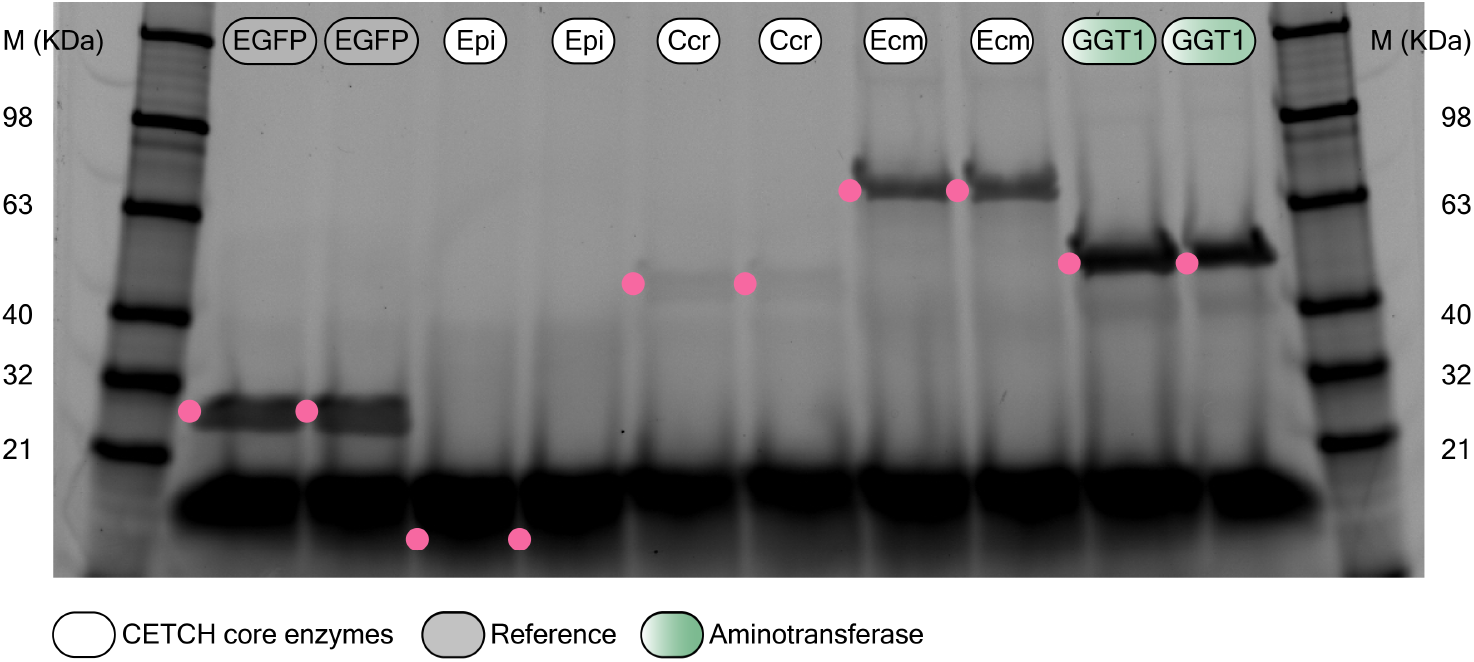
SDS PAGE of the proteins produced in PURE TX-TL system. The TX-TL of EGFP (27.8 kDa), Epi (16.8 kDa), Ccr (49.0 kDa), Ecm (73.7 kDa), GGT1 (55.8 kDa) was assessed by SDS–PAGE (without purification) on 4-20% gels, with FluoroTect GreenLys tRNA. Duplicates of each sample are analyzed; pink dots highlight sample bands.

**Fig. S9.**
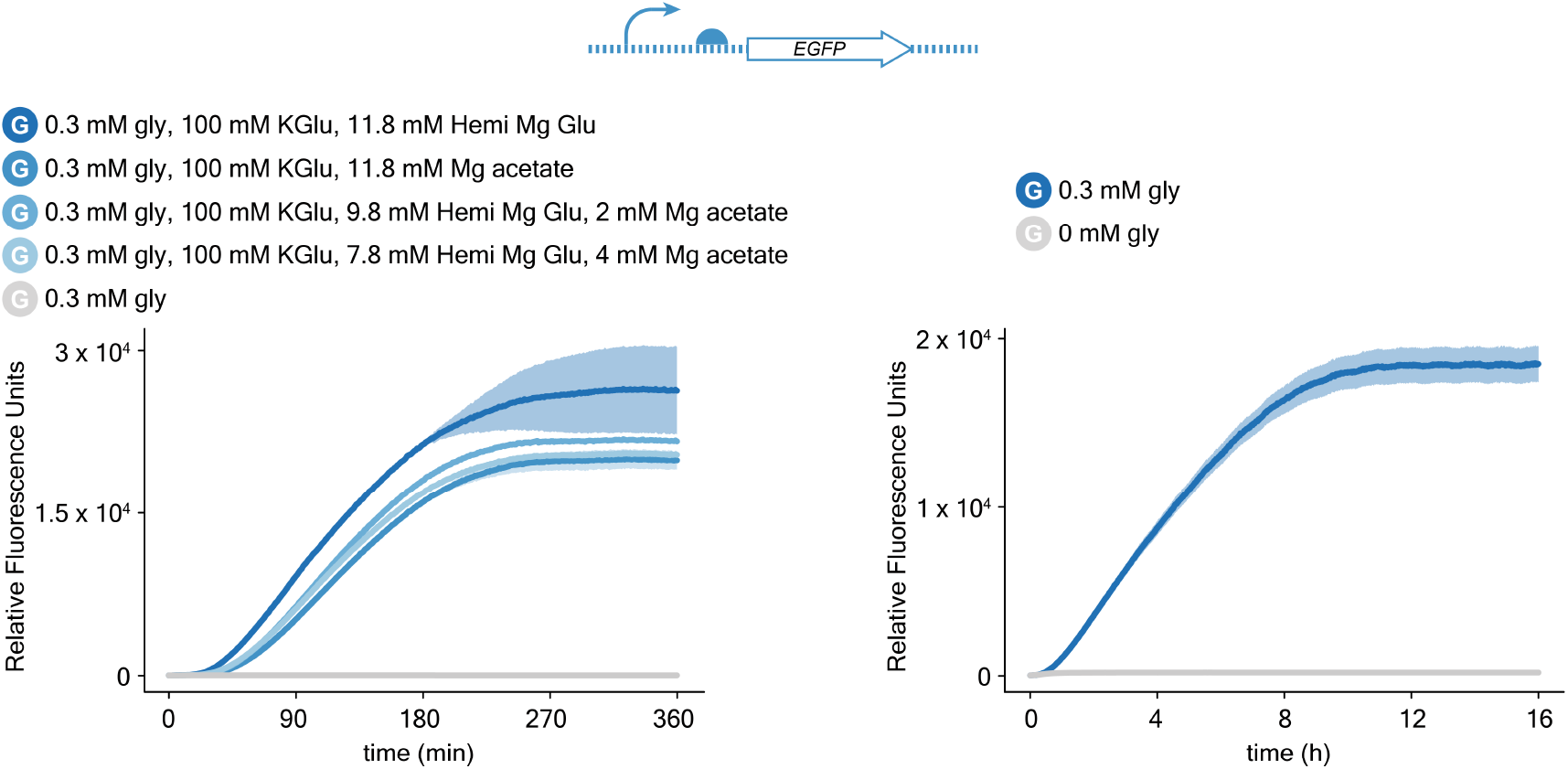
Cell-free TX-TL performed with different Mg^2+^ ion sources, or at lower temperature. Plots of the fluorescence signal resulting from the expression of *EGFP* template in a microplate reader. TX-TL reactions in buffer exchanged PURE system were supplemented with different sources of Mg^2+^ ion (left). A negative control without any source of Mg^2+^ was included (grey curve). TX-TL reactions in Δgly-PURE system were run at 30 °C for 16 h by supplementing the system with 300 μM glycine (right). A negative control without glycine was included (grey curve). TX–TL reactions (triplicates) were incubated into the microplate reader, with gain = 90. Curves represent the statistical mean of the results at any acquisition time, shadows represent the standard deviation of the same data.

**Fig. S10.**
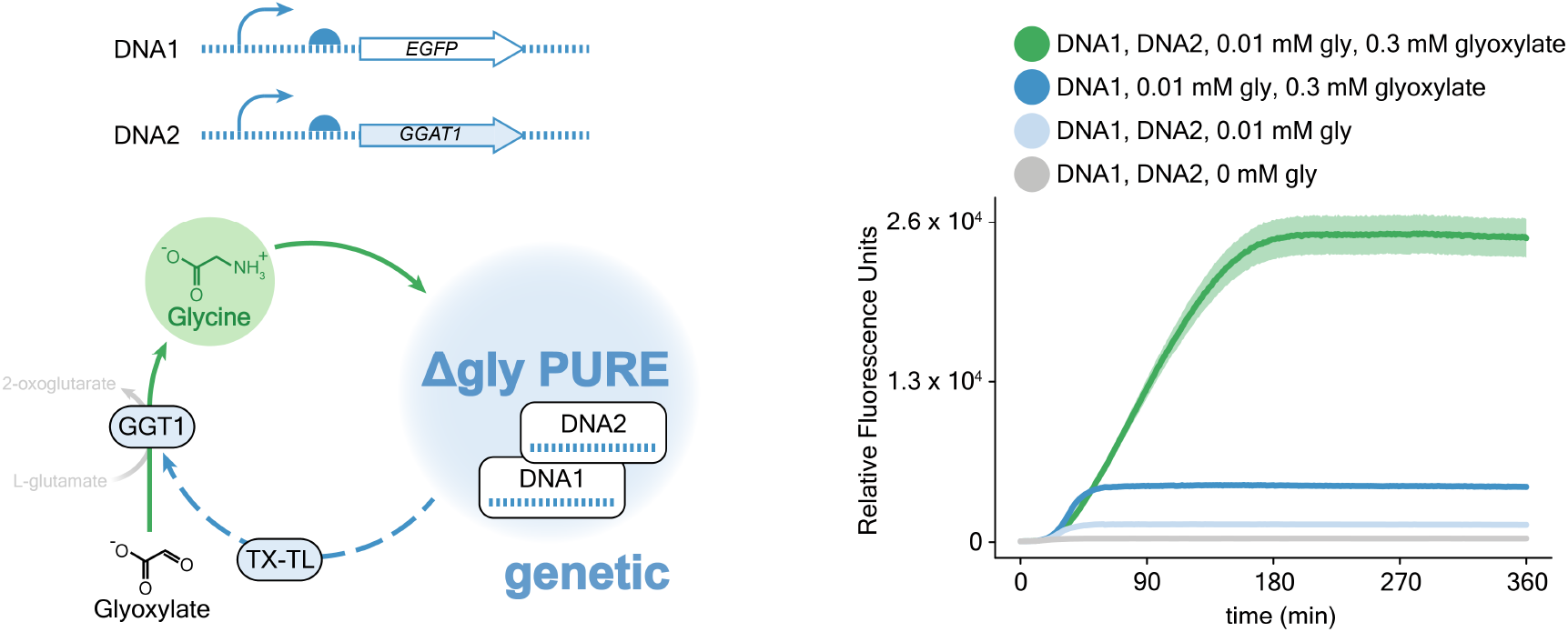
Activity assay for GGT1 produced by cell-free TX-TL. Sketch of the co-expression of *GGAT1* and *EGFP* DNA templates performed to assay the activity of the GGT1 enzyme produced by cell-free TX-TL in Δgly-PURE system (left). Plots of the fluorescence signal resulting from the expression of *EGFP* template in a microplate reader (right). The green curve is obtained by co-expressing *GGAT1* and *EGFP* templates in presence of 300 μM glyoxylate, and 10 μM glycine in a system missing the GGT1 enzyme. The light blue curve is obtained by co-expressing *Ecm* and *EGFP* templates in presence of 10 μM glycine. In the negative control experiment (grey curve) glycine is omitted. An additional control (dark blue curve) showing the result of the expression of the *EGFP* template in presence of 300 μM glyoxylate, and 10 μM glycine has been included. Reactions were incubated at 30 °C into the microplate reader for 6 h, with gain = 100. TX–TL reactions were run in triplicates. Curves represent the statistical mean of the results at any acquisition time, shadows represent the standard deviation of the same data.

**Fig. S11.**
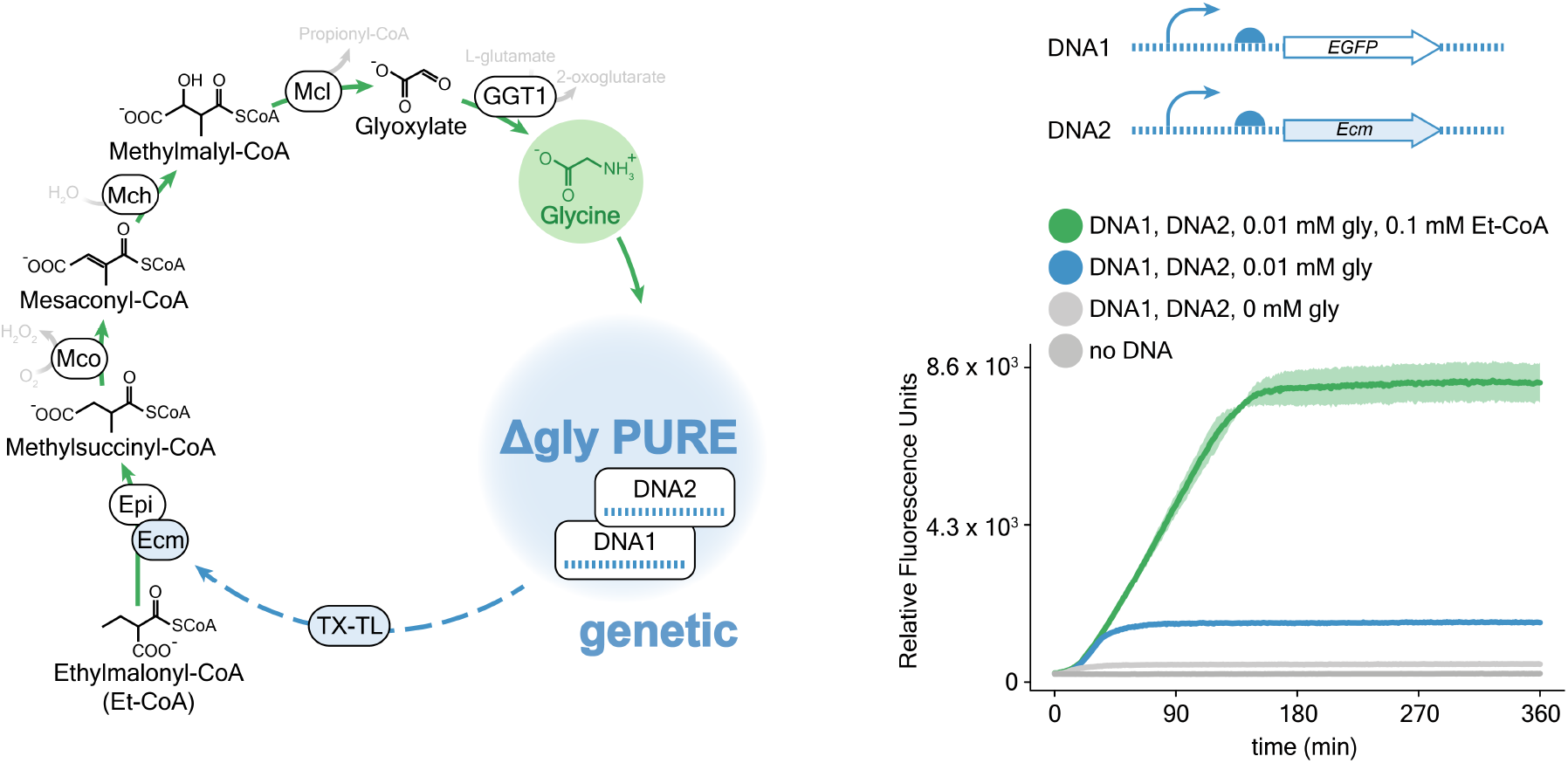
Activity assay for Ecm enzyme produced by cell-free TX-TL. Sketch of the co-expression of *Ecm* and *EGFP* DNA templates performed to assay the activity of the Ecm enzyme produced by cell-free TX-TL in Δgly-PURE system (left). Plots of the fluorescence signal resulting from the expression of *EGFP* template in a microplate reader (right). The green curve is obtained by co-expressing *Ecm* and *EGFP* templates in presence of 100 μM ethylmalonyl-CoA, and 10 μM glycine in a system missing the Ecm enzyme. The blue curve is obtained by co-expressing *Ecm* and *EGFP* templates in presence of 10 μM glycine. In the negative control experiments Δgly-PURE system is not provided with glycine (light grey curve), or DNA (dark grey curve). Reactions were incubated at 30 °C into the microplate reader for 6 h, with gain = 100. TX–TL reactions were run in duplicates. Curves represent the statistical mean of the results at any acquisition time, shadows represent the standard deviation of the same data.

**Fig. S12.**
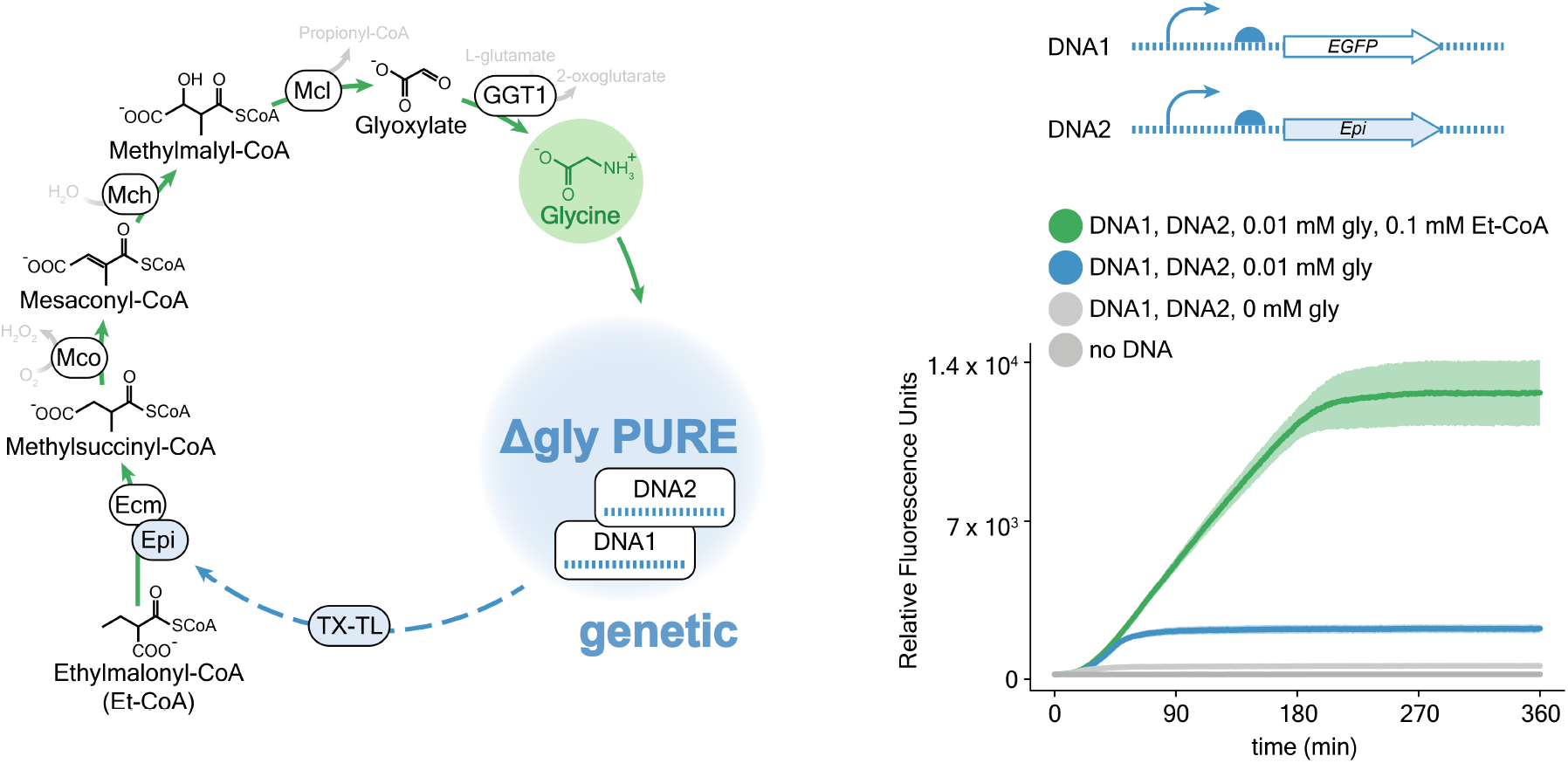
Activity assay for Epi enzyme produced by cell-free TX-TL. Sketch of the co-expression of *Epi* and *EGFP* DNA templates performed to assay the activity of the Epi enzyme produced by cell-free TX-TL in Δgly-PURE system (left). Plots of the fluorescence signal resulting from the expression of *EGFP* template in a microplate reader (right). The green curve is obtained by co-expressing *Epi* and *EGFP* templates in presence of 100 μM ethylmalonyl-CoA, and 10 μM glycine in a system missing the Epi enzyme. The blue curve is obtained by co-expressing *Epi* and *EGFP* templates in presence of 10 μM glycine. In the negative control experiments Δgly-PURE system is not provided with glycine (light grey curve), or DNA (dark grey curve). Reactions were incubated at 30 °C into the microplate reader for 6 h, with gain = 100. TX–TL reactions were run in triplicates. Curves represent the statistical mean of the results at any acquisition time, shadows represent the standard deviation of the same data.

**Fig. S13.**
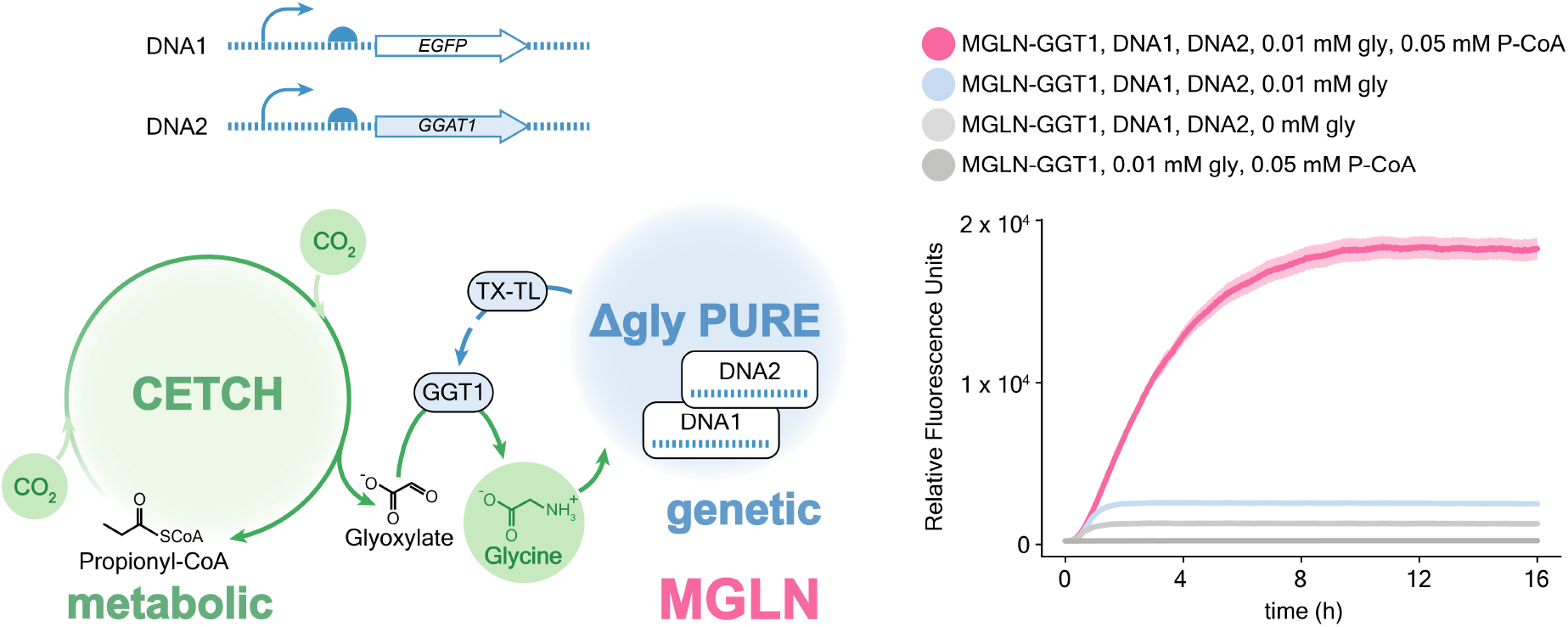
Autocatalytic MGLN. Sketch of a cell-free ΔGGT1-MGLN able to autonomously integrate its genetic (Δgly-PURE) and metabolic (CETCH cycle) layers in a feed forward fashion, by producing (TX-TL) the missing enzyme (GGT1) upon providing the system with low concentration of glycine (left). Plots of the fluorescence signal resulting from the expression of *EGFP* template in a microplate reader (right). The pink curve is obtained by supplementing the ΔGGT1-MGLN with *GGAT1* and *EGFP* templates, 50 μM propionyl-CoA, and 10 μM glycine. Negative controls without propionyl-CoA (blue curve), or without propionyl-CoA and glycine (light grey curve), or without DNA templates (dark grey curve) were performed. MGLN were incubated at 30 °C into the microplate reader for 16 h, with gain = 100. Reactions were run in triplicates. Curves represent the statistical mean of the results at any acquisition time, shadows represent the standard deviation of the same data.

**Fig. S14.**
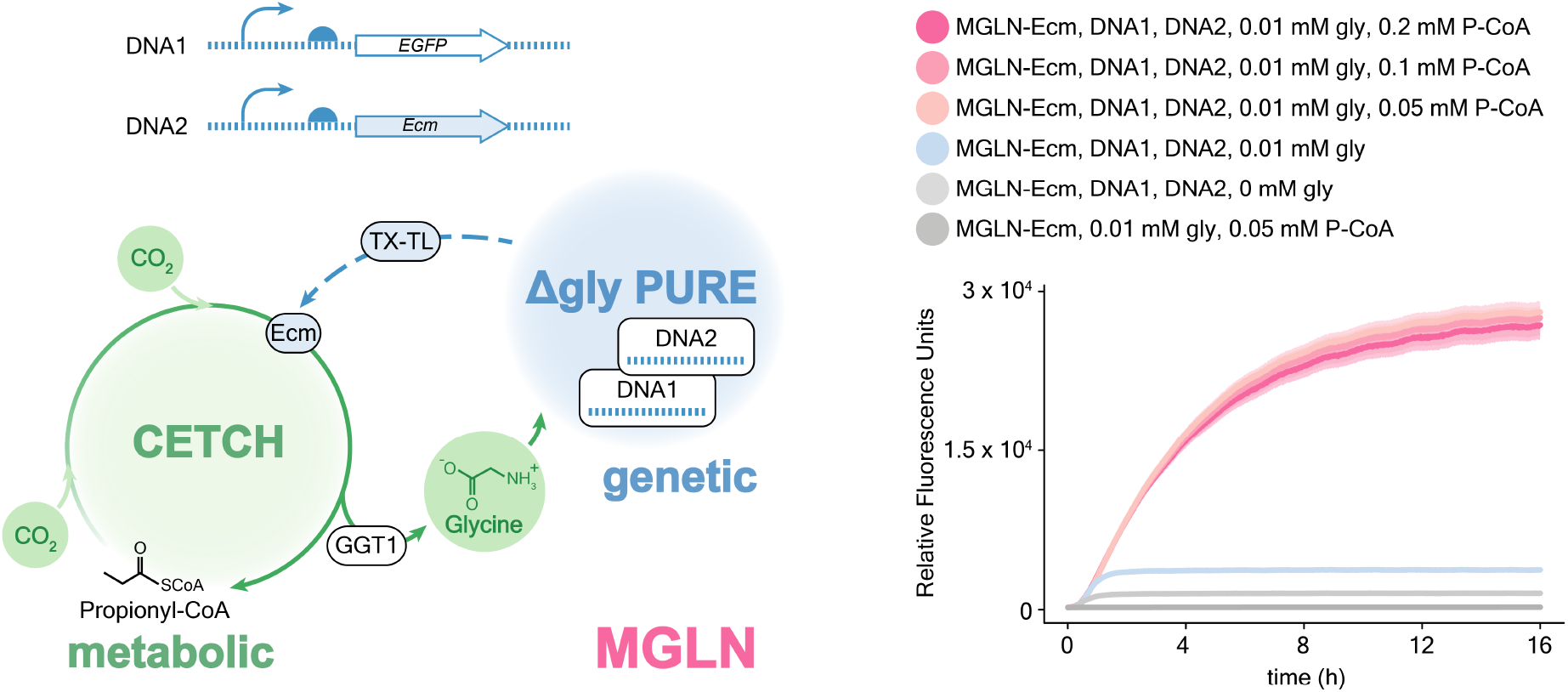
Self-regenerating MGLN. Sketch of a cell-free ΔEcm-MGLN able to partially self-regenerate (increase its CO_2_ fixation in a feed forward manner) by producing (TX-TL) the missing enzyme (Ecm) upon providing the system with low concentration of glycine (left). Plots of the fluorescence signal resulting from the expression of *EGFP* template in a microplate reader (right). The pink curves are obtained by supplementing the ΔEcm-MGLN with *Ecm* and *EGFP* templates, propionyl-CoA (200 μM, 100 μM, or 50 μM), and 10 μM glycine. Negative controls without propionyl-CoA (blue curve), or without propionyl-CoA and glycine (light grey curve), or without DNA templates (dark grey curve) were performed. MGLN were incubated at 30 °C into the microplate reader for 16 h, with gain = 100. Reactions were run in triplicates. Curves represent the statistical mean of the results at any acquisition time, shadows represent the standard deviation of the same data.

**Fig. S15.**
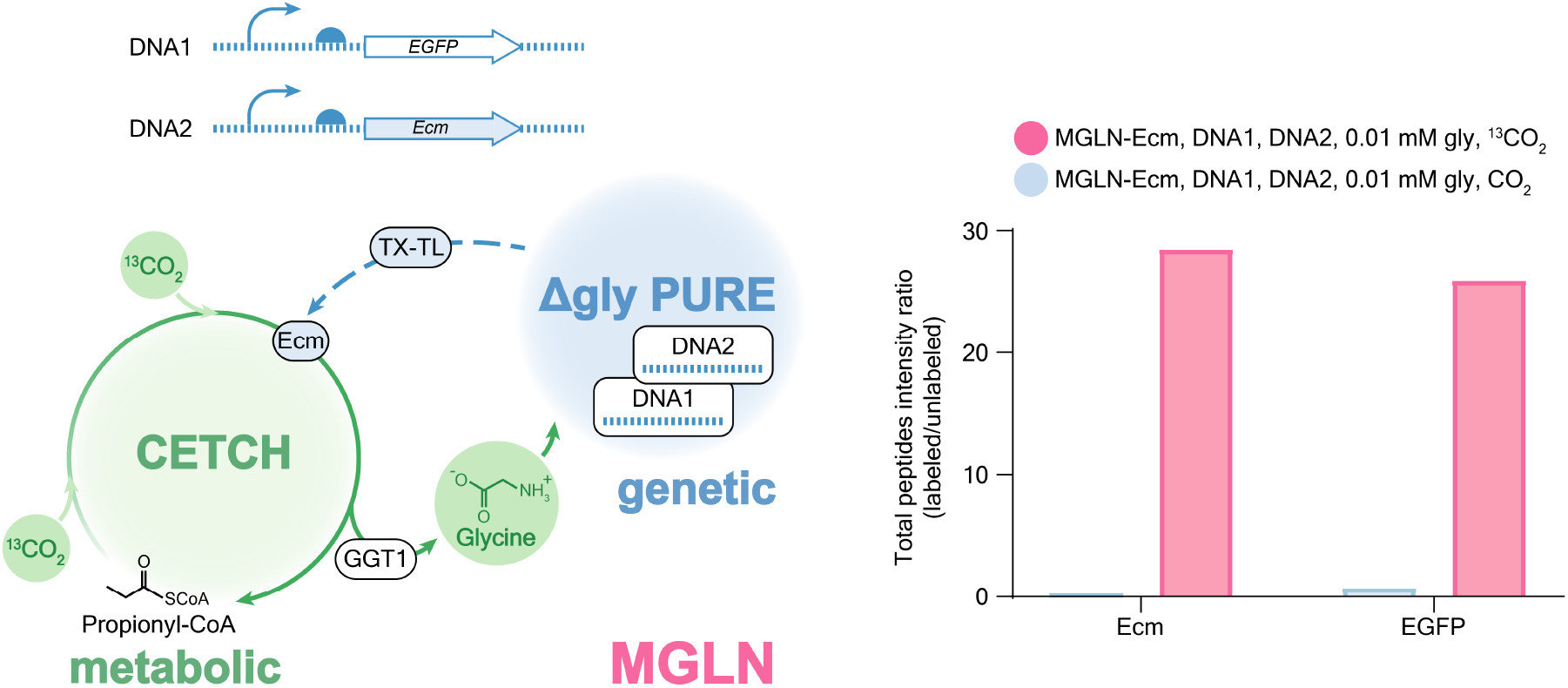
Self-regenerating MGLN: proteomic analysis. Sketch of a cell-free ΔEcm-MGLN able to partially self-regenerate (increase its CO_2_ fixation in a feed forward manner) by producing (TX-TL) the missing enzyme (Ecm) upon providing the system with low concentration of glycine (left). Plot of the ratio between the total intensity (sum) of the ^13^C-labeled peptides, and unlabeled peptides detected by proteomic analysis for Ecm, or EGFP, normalized by the labeled and unlabeled peptide intensity of most abundant protein (Mco) (right). Pink, or blue bars are obtained by supplementing the ΔEcm-MGLN with ^13^C-Sodium bicarbonate, or unlabeled Sodium bicarbonate respectively. MS raw data were analyzed with MaxQuant (version 2.0.3.0).

**Fig. S16.**
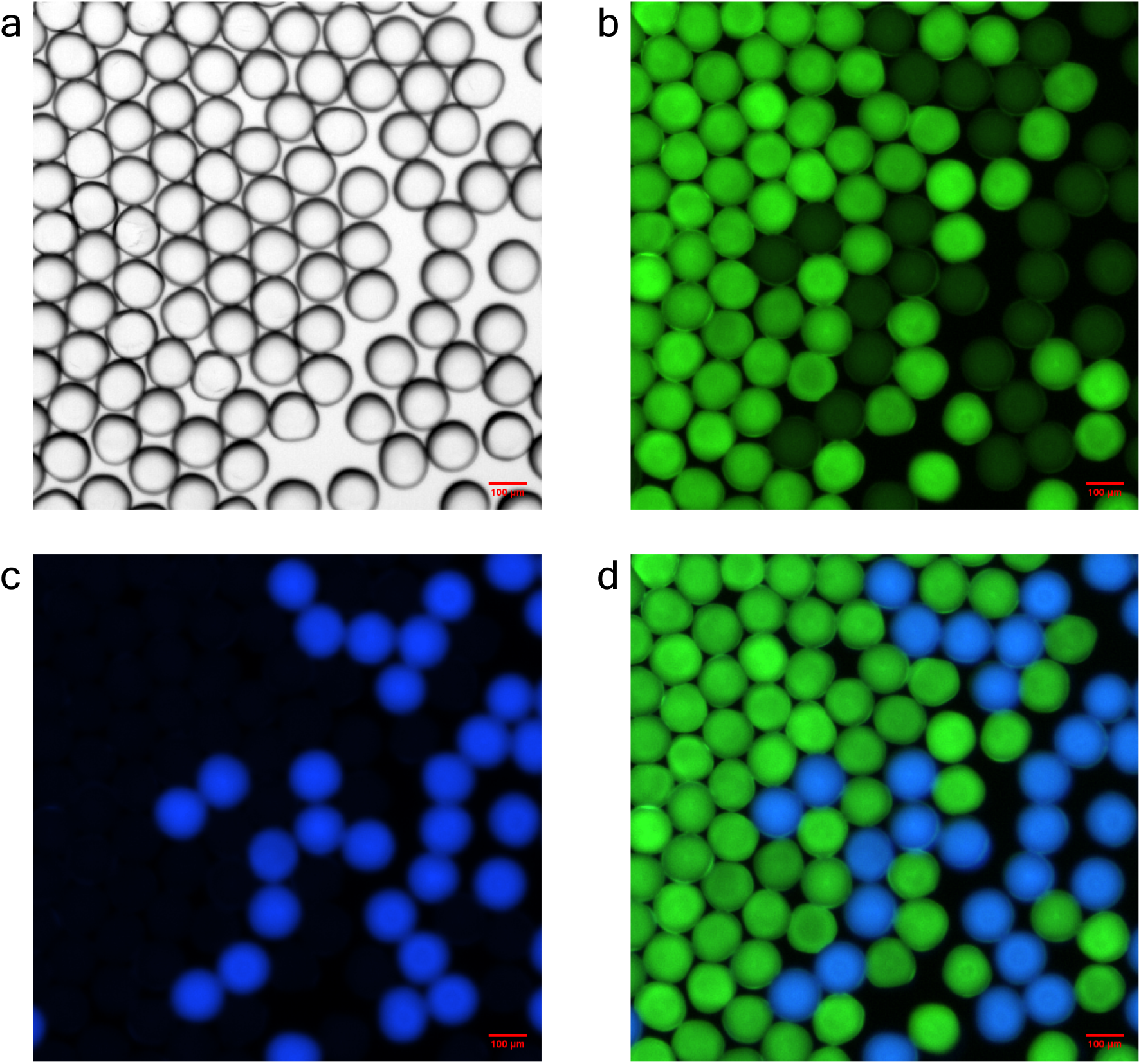
Self-regenerating MGLN compartmentalized in ∼100 μm droplets. Micrographs of a synthetic organelle-mimic able to increase its CO_2_ fixation autonomously in micrometer-size droplets, by bottom-up assembling a self-regenerating ΔEcm-MGLN. Droplets with *Ecm* and *EGFP* DNA templates, or without DNA (negative control) were incubated at 30 °C, and imaged after 6 h. The negative control droplets population was barcoded with Cascade Blue^TM^ hydrazide trilithium salt dye. (**a**) Brightfield image of the micrometer droplets; (**b**) sample droplet population (GFP channel); (**c**) negative control droplet population (DAPI channel); (**d**) overlay of the images acquired in GFP and DAPI channels.

**Table S1.** List of the enzymes overproduced in this study.

| Name | full name | origin | vector | plasmid | ref. |
| --- | --- | --- | --- | --- | --- |
| Pco | propionyl-CoA oxidase | *A. thaliana* | pET16b | pTE825 | (2) |
| Ccr | crotonyl-CoA carboxylase/reductase | *M. extorquens* | pET-23b(+) | pTE71 | (2) |
| Epi | methylmalonyl-/ethylmalonyl-CoA epimerase | *R. sphaeroides* | pET-16b | pTE45 | (2) |
| Mcm | methylmalonyl-CoA mutase | *R. sphaeroides* | pET-16b | pTE46 | (2) |
| Scr | succinyl-CoA reductase | *C. kluyveri* | pCDFDuet1 | pTE380 | (2) |
| Ssr | succinic semialdehyde reductase | *H. sapiens* | pLIC-SGC1 | P2BP1 | (2) |
| Hbs | 4-hydroxybutyryl-CoA synthetase | *N. maritimus* | pET-16b | pNMAR_0206 | (2) |
| Hbd | 4-hydroxybutyryl-CoA dehydratase | *N. maritimus* | pRSET-B | pTE393 | (2) |
| Ecm | ethylmalonyl-CoA mutase | *R. sphaeroides* | pET-16b | pTE33A | (2) |
| Mco | methylsuccinyl-CoA oxidase | *R. sphaeroides* | pET-16b | pTE813 | (2) |
| Mch | mesacony-CoA hydratase | *R. sphaeroides* | pET-16b | pMCH_Rs | (2) |
| Mcl1 | β-methylmalyl-CoA lyase | *R. sphaeroides* | pET-16b | pMCL1 Rs JZ 03 | (2) |
| KatE | catalase | *E. coli* | pCA24N | ASKA JW1721 | (2) |
| Fdh | formate dehydrogenase | *M. vaccae* | pET-21a | mutMycFDH | (2) |
| GGT1 | glutamate-glyoxylate aminotransferase1 | *A. thaliana* | pET-16b | pTE5153 | this work |
| BhcA | L-aspartate-glyoxylate aminotransferase | *P. denitrificans* | pET-16b | pTE538 | (8) |

**Table S2.** Kinetic parameters of the aminotransferase enzymes used in this study. Data are mean ± s.d., calculated by using GraphPad Prism 9, with nonlinear fits of 18 data points. Michaelis–Menten fits of enzyme kinetics, and SDS–PAGE gels showing purified proteins are provided in Fig. S1. (**) = measured in presence of 50 mM HEPES buffer pH 7.6, 11.8 mM Magnesium acetate, and 0.3 mM each of the 19 proteinogenic AAs-(glycine).

| Enzyme | substrates | $k_{cat}$<br>($s^{-1}$) | $K_M$<br>(mM) | $k_{cat} / K_M$<br>($mM^{-1} s^{-1}$) |
| --- | --- | --- | --- | --- |
| BhcA | glyoxylate and L-aspartate | $46 \pm 1.5$ | $0.51 \pm 0.06$ | $8.99 \times 10^4$ |
| | glyoxylate and L-glutamate | $11 \pm 1.5$ | $0.14 \pm 0.02$ | $8.14 \times 10^4$ |
| GGT1 | glyoxylate and L-glutamate | $87 \pm 2.4$ | $0.51 \pm 0.05$ | $17.2 \times 10^4$ |
| BhcA | glyoxylate and L-glutamate (**) | $10 \pm 0.5$ | $0.19 \pm 0.02$ | $5.14 \times 10^4$ |
| GGT1 | glyoxylate and L-glutamate (**) | $76 \pm 1.7$ | $0.33 \pm 0.03$ | $23.1 \times 10^4$ |

**Table S3.** MRM transitions of quantified residues (0.0 = unlabeled, 1.0 and 1.1 = ^15^N-labeled).

| Compound | precursor ion<br>MS1 (m/z) | product ion<br>MS2 (m/z) | fragmentor<br>(V) | collision<br>energy (V) | cell<br>accelerator (V) | polarity |
| --- | --- | --- | --- | --- | --- | --- |
| Ala 0.0 (quant) | 90 | 44.1 | 380 | 12 | 5 | Positive |
| Ala 1.0 (quant) | 91 | 44.1 | 380 | 12 | 5 | Positive |
| Ala 1.1 (quant) | 91 | 45.1 | 380 | 12 | 5 | Positive |
| Arg 0.0 (qual) | 174.9 | 116 | 380 | 12 | 5 | Positive |
| Arg 0.0 (quant) | 174.9 | 70.2 | 380 | 29 | 5 | Positive |
| Asn 0.0 (qual) | 133 | 87.1 | 380 | 17 | 5 | Positive |
| Asn 0.0 (quant) | 133 | 74.2 | 380 | 16 | 5 | Positive |
| Asn 1.0 (qual) | 134 | 87.1 | 380 | 17 | 5 | Positive |
| Asn 1.0 (quant) | 134 | 74.2 | 380 | 16 | 5 | Positive |
| Asn 1.1 (qual) | 134 | 88.1 | 380 | 17 | 5 | Positive |
| Asn 1.1 (quant) | 134 | 75.2 | 380 | 16 | 5 | Positive |
| Asp 0.0 (qual) | 134.1 | 88 | 380 | 9 | 5 | Positive |
| Asp 0.0 (quant) | 134.1 | 74 | 380 | 14 | 5 | Positive |
| Asp 1.0 (qual) | 135.1 | 88 | 380 | 9 | 5 | Positive |
| Asp 1.0 (quant) | 135.1 | 74 | 380 | 14 | 5 | Positive |
| Asp 1.1 (qual) | 135.1 | 89 | 380 | 9 | 5 | Positive |
| Asp 1.1 (quant) | 135.1 | 75 | 380 | 14 | 5 | Positive |
| Gln 0.0 (qual) | 147.2 | 130.1 | 380 | 8 | 5 | Positive |
| Gln 0.0 (quant) | 147.2 | 84.2 | 380 | 17 | 5 | Positive |
| Gly 0.0 (quant) | 76.1 | 30.3 | 380 | 32 | 5 | Positive |
| Gly 0.0 (qual) | 76.1 | 28.3 | 380 | 32 | 5 | Positive |
| Gly 1.0 (quant) | 77.1 | 30.3 | 380 | 32 | 5 | Positive |
| Gly 1.0 (qual) | 77.1 | 28.3 | 380 | 32 | 5 | Positive |
| Gly 1.1 (quant) | 77.1 | 31.3 | 380 | 32 | 5 | Positive |
| Gly 1.1 (qual) | 77.1 | 29.3 | 380 | 32 | 5 | Positive |
| His 0.0 (quant) | 156.1 | 110.1 | 380 | 16 | 5 | Positive |
| His 0.0 (qual) | 156.1 | 83 | 380 | 30 | 5 | Positive |
| Ile 0.0 (quant) | 132.1 | 86.1 | 380 | 8 | 5 | Positive |
| Ile 0.0 (qual) | 132.1 | 69.1 | 380 | 18 | 5 | Positive |
| Leu 0.0 (quant) | 132.1 | 86.1 | 380 | 8 | 5 | Positive |
| Leu 0.0 (qual) | 132.1 | 30.3 | 380 | 18 | 5 | Positive |
| Lys 0.0 (qual) | 147.1 | 130.1 | 380 | 8 | 5 | Positive |
| Lys 0.0 (quant) | 147.1 | 84.1 | 380 | 19 | 5 | Positive |
| Met 0.0 (qual) | 150.1 | 133 | 380 | 7 | 5 | Positive |
| Met 0.0 (quant) | 150.1 | 104 | 380 | 7 | 5 | Positive |
| Phe 0.0 (quant) | 166.1 | 120.2 | 380 | 13 | 5 | Positive |
| Phe 0.0 (qual) | 166.1 | 103.1 | 380 | 32 | 5 | Positive |
| Phe 1.0 (quant) | 167.1 | 120.2 | 380 | 13 | 5 | Positive |
| Phe 1.0 (qual) | 167.1 | 103.1 | 380 | 32 | 5 | Positive |
| Phe 1.1 (quant) | 167.1 | 121.2 | 380 | 13 | 5 | Positive |
| Pro 0.0 (quant) | 116 | 70 | 380 | 15 | 5 | Positive |
| Pro 0.0 (qual) | 116 | 43.3 | 380 | 35 | 5 | Positive |
| Ser 0.0 (quant) | 106.1 | 60.2 | 380 | 12 | 5 | Positive |
| Ser 0.0 (qual) | 106.1 | 42.2 | 380 | 11 | 5 | Positive |
| Ser 1.0 (quant) | 107.1 | 60.2 | 380 | 12 | 5 | Positive |
| Ser 1.0 (qual) | 107.1 | 42.2 | 380 | 11 | 5 | Positive |
| Ser 1.1 (quant) | 107.1 | 61.2 | 380 | 12 | 5 | Positive |
| Ser 1.1 (qual) | 107.1 | 43.2 | 380 | 11 | 5 | Positive |
| Thr 0.0 (quant) | 120.2 | 74.1 | 380 | 8 | 5 | Positive |
| Thr 0.0 (qual) | 120.2 | 55.9 | 380 | 18 | 5 | Positive |
| Trp 0.0 (quant) | 205.1 | 188 | 380 | 7 | 5 | Positive |
| Trp 0.0 (qual) | 205.1 | 145.9 | 380 | 17 | 5 | Positive |
| Trp 1.0 (quant) | 206.1 | 188 | 380 | 7 | 5 | Positive |
| Trp 1.0 (qual) | 206.1 | 145.9 | 380 | 17 | 5 | Positive |
| Trp 1.1 (quant) | 206.1 | 189 | 380 | 7 | 5 | Positive |
| Trp 1.1 (qual) | 206.1 | 146.9 | 380 | 17 | 5 | Positive |
| Tyr 0.0 (qual) | 182.1 | 165 | 380 | 6 | 5 | Positive |
| Tyr 0.0 (quant) | 182.1 | 136.1 | 380 | 12 | 5 | Positive |
| Tyr 1.0 (qual) | 183.1 | 165 | 380 | 6 | 5 | Positive |
| Tyr 1.0 (quant) | 183.1 | 136.1 | 380 | 12 | 5 | Positive |
| Tyr 1.1 (quant) | 183.1 | 137.1 | 380 | 12 | 5 | Positive |
| Val 0.0 (quant) | 118.1 | 72 | 380 | 9 | 5 | Positive |
| Val 0.0 (qual) | 118.1 | 55.1 | 380 | 23 | 5 | Positive |

**Table S4.** MRM transitions of ^13^C2-glycine internal standard (IS).

| Compound | precursor ion<br>MS1 (m/z) | product ion<br>MS2 (m/z) | fragmentor<br>(V) | collision<br>energy (V) | cell<br>accelerator (V) | polarity |
| --- | --- | --- | --- | --- | --- | --- |
| Gly IS (qual) | 78.1 | 32.3 | 380 | 32 | 5 | Positive |
| Gly IS (quant) | 78.1 | 31.3 | 380 | 32 | 5 | Positive |
| Gly IS (qual) | 78.1 | 30.3 | 380 | 32 | 5 | Positive |
| Gly IS (qual) | 78.1 | 29.3 | 380 | 32 | 5 | Positive |
| Gly IS (qual) | 78.1 | 28.3 | 380 | 32 | 5 | Positive |

**Table S5.** Detailed composition of CETCH assays (1–10) stepwise supplemented with PURE TX-TL components. (b.e. = buffer exchanged).

| CETCH components | (1) | (2) | (3) | (4) | (5) | (6) | (7) | (8) | (9) | (10) |
| --- | --- | --- | --- | --- | --- | --- | --- | --- | --- | --- |
| μM Pco | 0.5 | 0.5 | 0.5 | 0.5 | 0.5 | 0.5 | 0.5 | 0.5 | 0.5 | 0.5 |
| μM Ccr | 0.4 | 0.4 | 0.4 | 0.4 | 0.4 | 0.4 | 0.4 | 0.4 | 0.4 | 0.4 |
| μM Epi | 0.3 | 0.3 | 0.3 | 0.3 | 0.3 | 0.3 | 0.3 | 0.3 | 0.3 | 0.3 |
| μM Mcm | 0.3 | 0.3 | 0.3 | 0.3 | 0.3 | 0.3 | 0.3 | 0.3 | 0.3 | 0.3 |
| μM Scr | 1.4 | 1.4 | 1.4 | 1.4 | 1.4 | 1.4 | 1.4 | 1.4 | 1.4 | 1.4 |
| μM Ssr | 0.9 | 0.9 | 0.9 | 0.9 | 0.9 | 0.9 | 0.9 | 0.9 | 0.9 | 0.9 |
| μM Hbs | 0.1 | 0.1 | 0.1 | 0.1 | 0.1 | 0.1 | 0.1 | 0.1 | 0.1 | 0.1 |
| μM Hbd | 0.3 | 0.3 | 0.3 | 0.3 | 0.3 | 0.3 | 0.3 | 0.3 | 0.3 | 0.3 |
| μM Ecm | 0.6 | 0.6 | 0.6 | 0.6 | 0.6 | 0.6 | 0.6 | 0.6 | 0.6 | 0.6 |
| μM Mco | 9.3 | 9.3 | 9.3 | 9.3 | 9.3 | 9.3 | 9.3 | 9.3 | 9.3 | 9.3 |
| μM Mch | 0.1 | 0.1 | 0.1 | 0.1 | 0.1 | 0.1 | 0.1 | 0.1 | 0.1 | 0.1 |
| μM Mcl1 | 2.9 | 2.9 | 2.9 | 2.9 | 2.9 | 2.9 | 2.9 | 2.9 | 2.9 | 2.9 |
| μM KatE | 0.3 | 0.3 | 0.3 | 0.3 | 0.3 | 0.3 | 0.3 | 0.3 | 0.3 | 0.3 |
| μM Fdh | 2.6 | 2.6 | 2.6 | 2.6 | 2.6 | 2.6 | 2.6 | 2.6 | 2.6 | 2.6 |
| μM CK | 0.5 | 0.5 | 0.5 | 0.5 | 0.5 | 0.5 | 0.5 | 0.5 | 0.5 | 0.5 |
| nM CA | 6 | 6 | 6 | 6 | 6 | 6 | 6 | 6 | 6 | 6 |
| nM GGT1 | 20 | 20 | 20 | 20 | 20 | 20 | 20 | 20 | 20 | 20 |
| mM HEPES pH 7.8 | 75 | 75 | 75 | 75 | 75 | 75 | 75 | 75 | 75 | 75 |
| mM MgCl<sub>2</sub> | 2.5 | 2.5 | 2.5 | 2.5 | 2.5 | 2.5 | 2.5 | 2.5 | 2.5 | 2.5 |
| mM CP | 1 | 1 | 1 | 1 | 1 | 1 | 1 | 21 | 21 | 21 |
| mM NaHCO<sub>3</sub> | 2 | 2 | 2 | 2 | 2 | 2 | 2 | 2 | 2 | 2 |
| mM HCOONa | 8 | 8 | 8 | 8 | 8 | 8 | 8 | 8 | 8 | 8 |
| mM CoA | 0.1 | 0.1 | 0.1 | 0.1 | 0.1 | 0.1 | 0.1 | 0.1 | 0.1 | 0.1 |
| mM ATP | 2 | 2 | 2 | 2 | 4 | 2 | 2 | 2 | 4 | 4 |
| mM NADPH | 0.75 | 0.75 | 0.75 | 0.75 | 0.75 | 0.75 | 0.75 | 0.75 | 0.75 | 0.75 |
| μM PLP | 20 | 20 | 20 | 20 | 20 | 20 | 20 | 20 | 20 | 20 |
| mM KGlu | 100 | 100 | 100 | 100 | 100 | 100 | 100 | 100 | 100 | 100 |
| mM GTP | - | - | - | - | 2 | - | - | - | 2 | 2 |
| mM CTP | - | - | - | - | 1 | - | - | - | 1 | 1 |
| mM UTP | - | - | - | - | 1 | - | - | - | 1 | 1 |
| mM DTT | - | 1 | 1 | 1 | 1 | 1 | 1 | 1 | 1 | 1 |
| mM TCEP | - | - | 1 | - | - | - | - | - | - | - |
| mM Mg acetate | - | - | - | 11.8 | - | - | - | - | - | - |
| mM hemi Mg glu | - | - | - | - | 11.8 | 11.8 | 11.8 | 11.8 | 11.8 | 11.8 |
| mM 19 AAs (each) | - | - | - | - | 0.3 | 0.3 | 0.3 | 0.3 | 0.3 | 0.3 |
| μM folinic acid | - | - | - | - | - | 20 | - | - | 20 | 20 |
| mM spermidine | - | - | - | - | - | - | 2 | - | 2 | 2 |
| mM propionyl-CoA | 0.2 | 0.2 | 0.2 | 0.2 | 0.2 | 0.2 | 0.2 | 0.2 | 0.2 | 0.2 |
| v/v RNase inhibitor | - | - | - | - | 1/49 | 1/49 | 1/49 | 1/49 | 1/49 | 1/49 |
| v/v TX-TL enzymes | - | - | - | - | 1/19 | 1/19 | 1/19 | 1/19 | 1/19 | - |
| v/v b.e. TX-TL enzymes | - | - | - | - | - | - | - | - | - | 1/19 |
| v/v ribosomes | - | - | - | - | 1/19 | 1/19 | 1/19 | 1/19 | 1/19 | - |
| v/v b.e. ribosomes | - | - | - | - | - | - | - | - | - | 1/19 |

**Table S6.**
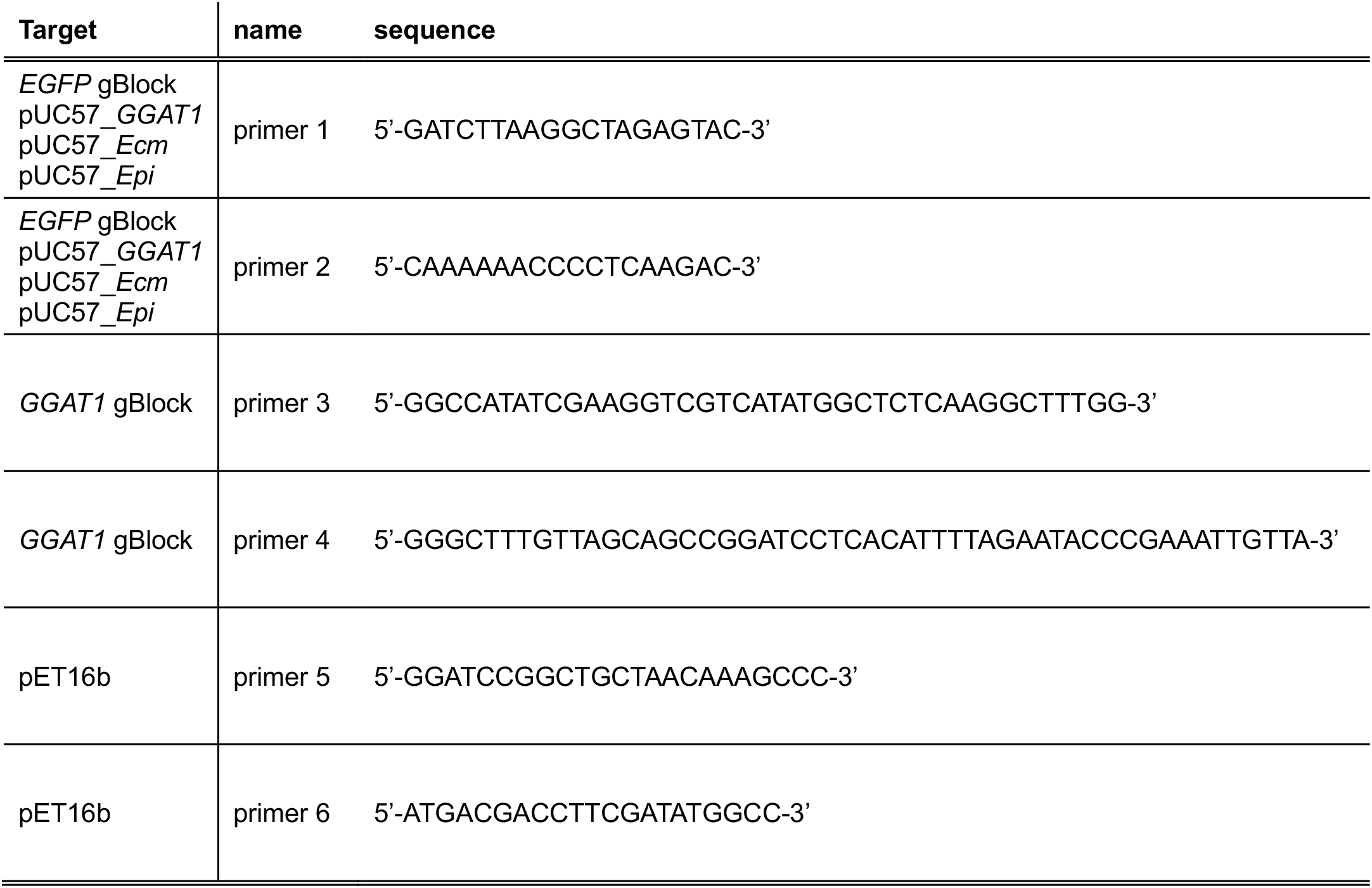
Primers used in this study.

**Table S7.**
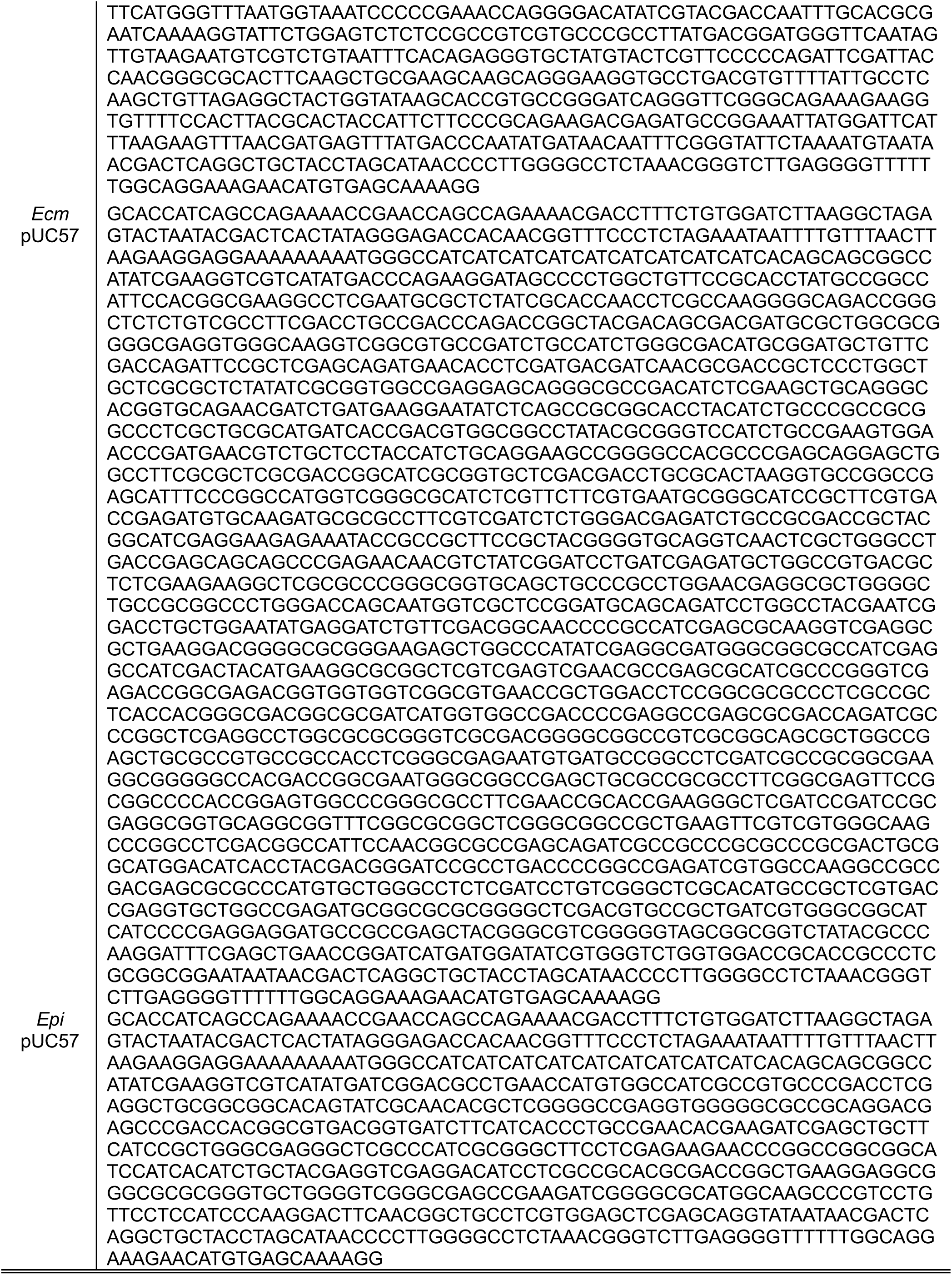
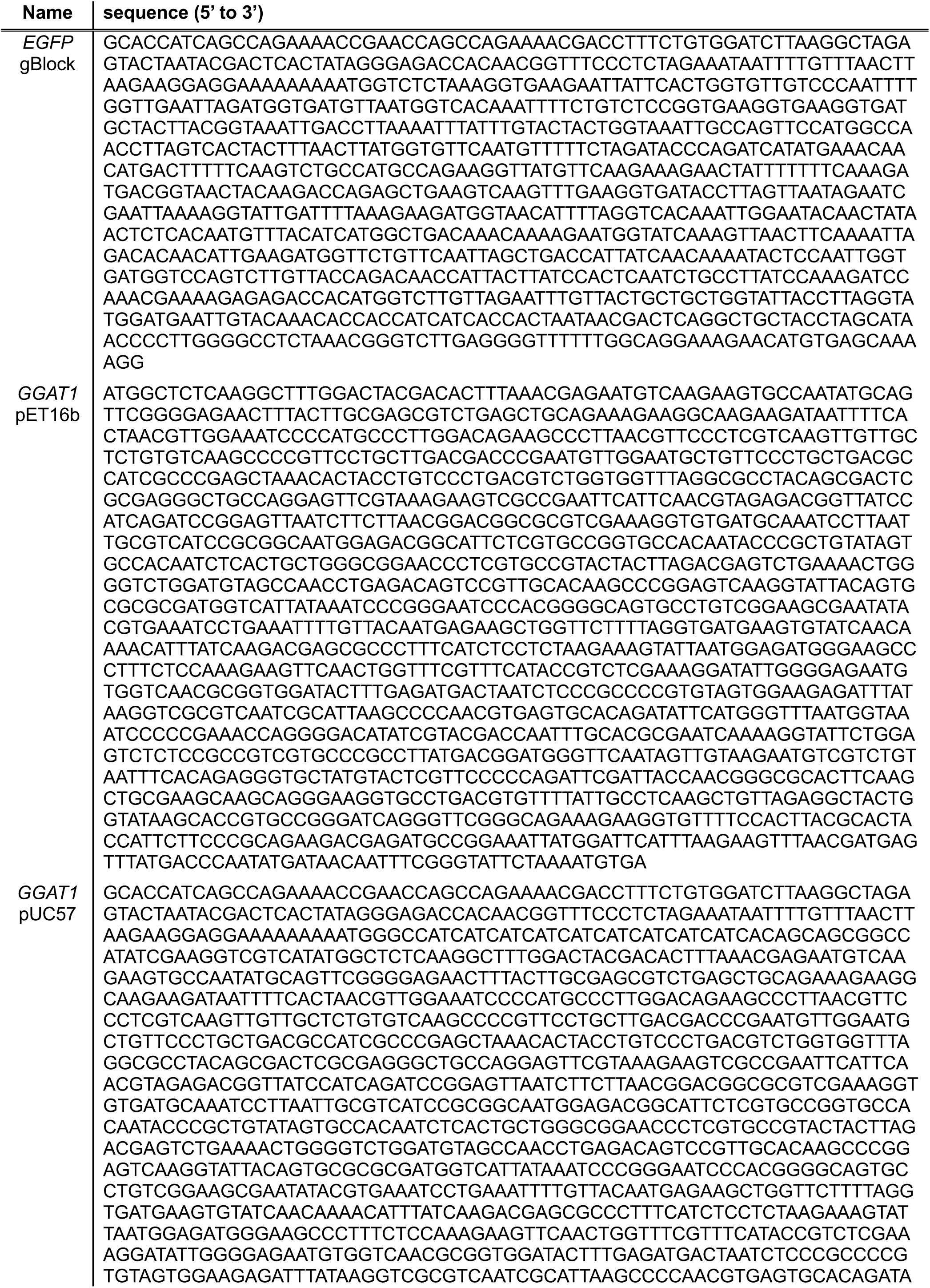
DNA sequences of the genes used in this study.

## Notes

### Competing Interest Statement

The authors have declared no competing interest.

